# Multimodal phenotyping reveals plant-food-related dietary information in serum Raman spectra of community-dwelling Japanese adults

**DOI:** 10.64898/2026.10.08.754882

**Authors:** Kenichi Hiraga, Takafumi Miyamoto, Teppei Nishino, Jieun Yoon, Kyohei Shibuya, Xiucai Ye, Tianyi Shi, Weike Liu, Hisato Suzuki, Hiroko Isoda, Tetsuya Sakurai, Hiroyuki Nishiyama, Tomohiro Okura, Motohiro Sekiya

**Affiliations:** Center for Cyber Medicine Research, University of Tsukuba, Tsukuba, Ibaraki 305-8575, Japan; Organization for Human Smart City Connection, University of Tsukuba, Tsukuba, Ibaraki 305-8577, Japan; Department of Clinical Medicine, Institute of Medicine, University of Tsukuba, Tsukuba, Ibaraki 305-8575, Japan; Department of Endocrinology and Metabolism, Institute of Medicine, University of Tsukuba, Tsukuba, Ibaraki 305-8575, Japan; Transborder Medical Research Center, University of Tsukuba, Tsukuba, Ibaraki 305-8575, Japan; Institute of Health and Sport Sciences, University of Tsukuba, Ibaraki 305-8574, Japan; R&D Center for Tailor-Made QOL, University of Tsukuba, Ibaraki 305-8550, Japan; Department of Computer Science, University of Tsukuba, Tsukuba, Ibaraki 305-8577, Japan; Tsukuba Life Science Innovation Program, University of Tsukuba, Tsukuba 305-8577, Japan; Center for Medical Genetics, Keio University School of Medicine, Tokyo 160-8582, Japan; Institute of Human Sciences, University of Tsukuba, Tsukuba, Ibaraki 305-8572, Japan; Institute of Life and Environmental Sciences, University of Tsukuba, Tsukuba, Ibaraki 305-8572, Japan; Department of Urology, University of Tsukuba, Tsukuba, Ibaraki 305-8575, Japan

## Abstract

Healthy dietary patterns are associated with long-term health, but diet-quality scores do not reveal how habitual dietary information is represented in human biological phenotypes. Here we linked habitual dietary assessment with preceding health records and contemporaneous clinical chemistry, serum metabolomics and serum Raman spectra in community-dwelling Japanese adults. Higher diet quality corresponded to a more favorable preceding health profile, but did not define discrete states across contemporary multivariate phenotypes. Instead, specific dietary information was recoverable across these layers: Raman spectra provided predictive information complementary to clinical and metabolomic measurements for multiple dietary variables. Raman variation corresponding to β-carotene equivalent could be traced to carotenoid-assigned bands and aligned with a broader plant-food-related dietary dimension. These findings show that habitual diet is represented not as a single biological state but through biologically traceable dietary dimensions distributed across complementary phenotypic layers.

## Introduction

Diet is one of the major modifiable determinants of human health. The Alternative Healthy Eating Index 2010 (AHEI-2010), widely used to assess overall diet quality, integrates multiple foods and nutrients on the basis of their associations with chronic disease risk, and higher AHEI-2010 scores are associated with lower risks of major chronic diseases, including coronary heart disease and type 2 diabetes^1^. Long-term maintenance of healthy dietary patterns is also associated with healthy ageing^2^, and studies in Japanese populations have reported associations between higher diet quality and lower all-cause or cardiovascular mortality^3^K^4^. Together, these findings underscore the importance of evaluating diet not only in terms of individual nutrients but also as an integrated pattern composed of multiple foods and nutrients.

Diet-quality scores, however, summarize an individual’s diet into a single value and do not directly indicate how the underlying dietary information is represented in that individual’s biological state. Individuals with similar diet-quality scores may achieve those scores through different combinations of foods and nutrients. Moreover, even when individuals consume the same or similar foods, metabolic responses such as postprandial glucose, insulin and triglycerides vary substantially among individuals^5^K^6^. Understanding the relationship between diet and health at the individual level therefore requires not only assessment of dietary exposure but also identification of which aspects of diet are manifested in which molecular and physiological phenotypes. This challenge may be particularly relevant in generally healthy populations, in whom diet-related biological differences are likely to be subtle and may be distributed across multiple phenotypic dimensions rather than expressed as clearly separable biological states.

Diet-related biological variation can be detected across several measurement modalities. Routine blood tests have been associated with healthy dietary patterns across multiple clinical and biochemical markers^7^, and circulating metabolomic profiles associated with healthy dietary patterns, including AHEI-2010, have also been identified^8^K^9^. Raman spectroscopy provides label-free information on the molecular composition of biological specimens and has been applied to detect molecular information related to diet and nutritional status^10^K^11^. Together, these observations indicate that dietary information can be reflected across different biological phenotype layers. However, it remains unclear, within the same individuals, how overall diet quality and more specific food- and nutrient-related information are distributed across, or captured complementarily by, these layers, particularly when such differences are not large enough to define clearly distinct phenotypic states.

Here, in community-dwelling Japanese adults from the Tsukuba Happiness Life Study^12–14^, we linked habitual dietary assessment at the individual level to temporally preceding health-insurance claims and municipal health-check information, as well as to contemporaneously acquired blood and urine tests, serum metabolomics and serum Raman spectra. We examined how overall diet quality corresponded to preceding health background and contemporary multidimensional biological phenotypes, and how more specific diet-related information was represented across these phenotypic layers. Higher diet quality corresponded to a more favorable preceding health profile, whereas diet-quality quartiles did not separate contemporary multidimensional phenotypes into clearly discrete states. By contrast, when we examined more specific dietary information, Raman spectra provided additional predictive information beyond blood tests and serum metabolomics for multiple dietary variables. Raman variation corresponding to β-carotene equivalent could be traced to established carotenoid-assigned bands and aligned with a broader plant-food-related dietary dimension. Together, these findings suggest that habitual diet is represented not as a single biological state but through biologically traceable dietary dimensions distributed across complementary phenotypic layers.

## Results

### Study overview and analytical populations

In FY2022 of the Tsukuba Happiness Life Study, participants underwent dietary assessment, general health examination, blood and urine testing, serum metabolomics and serum Raman measurements (Fig. 1). Of the 405 participants who attended the health examination, 401 had available Brief-type Self-administered Diet History Questionnaire (BDHQ) data^15^K^16^. The 367 participants with complete blood and urine tests, serum metabolomics, Raman spectra and BDHQ data were included in the integrated analyses of multilayer phenotypes and diet. To assess preceding health status, we additionally linked these data to municipal records held by Tsukuba City from 2014–2018. Among the 401 participants with BDHQ data, at least one municipal record during the observation period was identified for 132 participants; at least one medical claim was identified for 129 participants, and municipal health-check measurements were available for 93 participants.

**Fig. 1.**
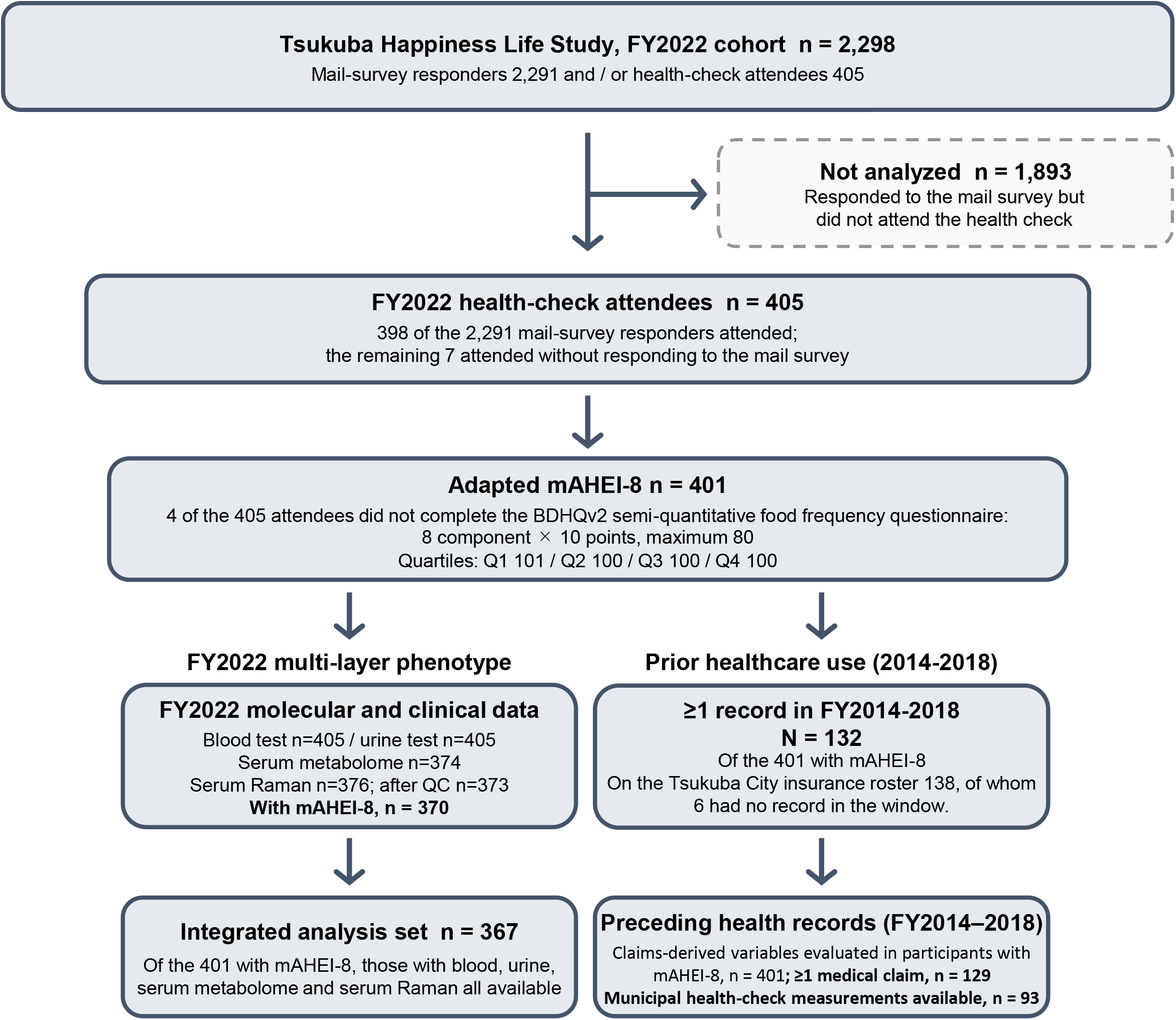
Analytical populations in the FY2022 Tsukuba Happiness Life Study cohort. Participant flow from the FY2022 cohort (n = 2,298) to the principal analytical populations. Among the 405 health-examination participants, BDHQ data were available for 401, for whom mAHEI-8 could be calculated. Blood and urine tests were available for 405 participants, serum metabolomics for 374 and serum Raman spectra for 376; three spectra were excluded during technical quality control, leaving 373 for Raman analyses. The integrated diet–multilayer phenotype analyses included 367 participants with available BDHQ, blood/urine, serum metabolome and quality-controlled Raman data. Among the 401 participants with mAHEI-8, at least one municipal record from 2014–2018 was identified for 132 participants; medical claims were identified for 129 and municipal health-check measurements for 93.

### mAHEI-8 captured a heterogeneous diet-quality gradient

Habitual diet quality in this population was assessed using an 8-component modified AHEI-2010 score (mAHEI-8; range 0–80), based on AHEI-2010 and comprising eight components that could be consistently evaluated from the BDHQ: vegetables, fruit, legumes, EPA + DHA, polyunsaturated fatty acids (PUFA), sugar-sweetened beverages + fruit juice, red + processed meat and sodium (Supplementary Table 1). When BDHQ-derived intakes were converted to component scores of 0–10 according to AHEI-2010 scoring rules, the score distributions differed substantially across components (Supplementary Fig. 1a,b). For EPA + DHA, 93.8% of participants reached the maximum score, whereas fruit had the lowest mean score among the eight components (3.11 points). Sodium was scored by population intake deciles, as in AHEI-2010; because sodium intake differed clearly by sex in this population (4,768 mg per day in men versus 4,157 mg per day in women, P = 2.8 × 10^−6^), deciles were defined within sex (Supplementary Fig. 1c).

In FY2022 of the Tsukuba Happiness Life Study, mAHEI-8 could be calculated for 401 participants, who were classified into quartiles according to its distribution (Q1, n = 101; Q2–Q4, n = 100 each; Fig. 2a). Mean mAHEI-8 scores across Q1–Q4 were 40.7 ± 3.0, 46.5 ± 1.4, 51.1 ± 1.3 and 57.1 ± 3.0, respectively; the overall mean was 48.9 ± 6.5 (range, 30.7–70.7). Mean age was nearly constant across Q1–Q4 (65.3, 65.0, 65.6 and 65.4 years), with no trend across quartiles (Fig. 2b; P for trend = 0.802; omnibus P = 0.973). By contrast, BMI tended to decrease across higher mAHEI-8 quartiles (23.7, 22.9, 23.1 and 22.4 kg m⁻²; Fig. 2b; P for trend = 0.006; omnibus P = 0.021). Total energy intake across Q1–Q4 was 1,881, 1,974, 1,956 and 1,752 kcal per day, respectively, and was lowest in Q4, although the pattern was not monotonic (Fig. 2b; P for trend = 0.085; omnibus P = 0.017). The proportions of women and participants who had never smoked were higher in the higher-mAHEI-8 groups, whereas educational attainment and perceived economic status showed no clear differences (Fig. 2c). Thus, mAHEI-8 quartiles in this population represented a diet-quality gradient accompanied by differences in selected lifestyle and anthropometric characteristics, rather than a uniform gradient of age or socioeconomic background.

**Fig. 2.**
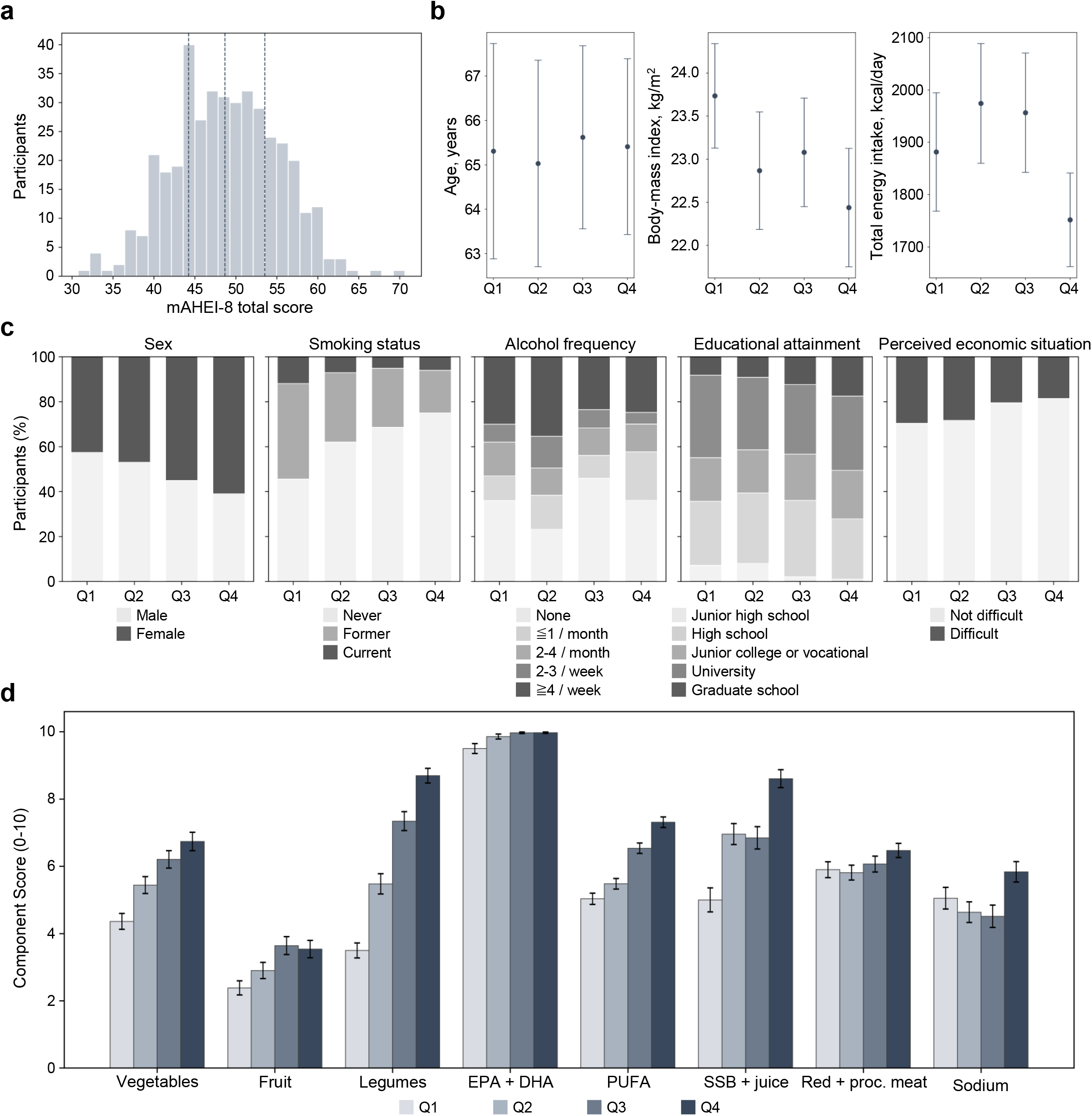
The mAHEI-8 gradient and its composition. The analytical population comprised 401 participants for whom mAHEI-8 could be calculated (Q1, n = 101; Q2–Q4, n = 100 each). a, Distribution of total mAHEI-8 scores and quartile boundaries. b, Quartile-specific means and 95% confidence intervals for age, BMI and total energy intake; BMI was available for 391 participants. P values are from Spearman tests for trend across ordinal quartiles and Kruskal–Wallis tests for overall differences. c, Quartile-specific proportions of participant characteristics and χ² tests. d, Mean component scores ± s.e.m. across quartiles for the eight components.

To identify the dietary components underlying this gradient, we decomposed the 16.4-point difference in mean mAHEI-8 between Q1 and Q4 across the eight components. Legumes (+5.2 points), sugar-sweetened beverages + fruit juice (+3.6 points), vegetables (+2.4 points) and PUFA (+2.3 points) together accounted for approximately 82% of the total score difference (Fig. 2d). By contrast, 93.8% of participants received the maximum EPA + DHA score, resulting in little contribution of this component to quartile separation. The mAHEI-8 gradient in this population was therefore not generated by uniform changes across all eight components, but was characterized primarily by differences in legumes, sugar-sweetened beverages + fruit juice, vegetables and PUFA.

### Higher mAHEI-8 corresponded to a more favorable preceding health profile

Healthy dietary patterns, including AHEI-2010, have been associated with lower chronic disease risk and healthier ageing¹K². We therefore asked whether the mAHEI-8 gradient observed in this population also corresponded to a longer-term health background. The proportions of participants with at least one municipal record from 2014–2018 were 31.7%, 41.0%, 31.0% and 28.0% across Q1–Q4, respectively, with no significant difference among quartiles (χ² P = 0.232; Supplementary Fig. 2a). Participants with a municipal record were, however, older than those without one (73.3 versus 61.4 years, P = 1.5 × 10⁻²⁴; Supplementary Fig. 2b).

An overview of preceding health information by quartile showed that the higher-mAHEI-8 groups tended to have a lower prevalence of hospitalization and fewer total medical-care days. Among the available historical health-check measures, multiple indicators also generally shifted in a more favorable direction, including lower BMI, waist circumference, systolic blood pressure, LDL cholesterol, triglycerides and creatinine, and higher eGFR (Fig. 3a). Not all indicators changed monotonically; nevertheless, similar directional patterns for several measures remained evident after stratification by age and sex, particularly among participants aged ≥65 years (Supplementary Fig. 3).

**Fig. 3.**
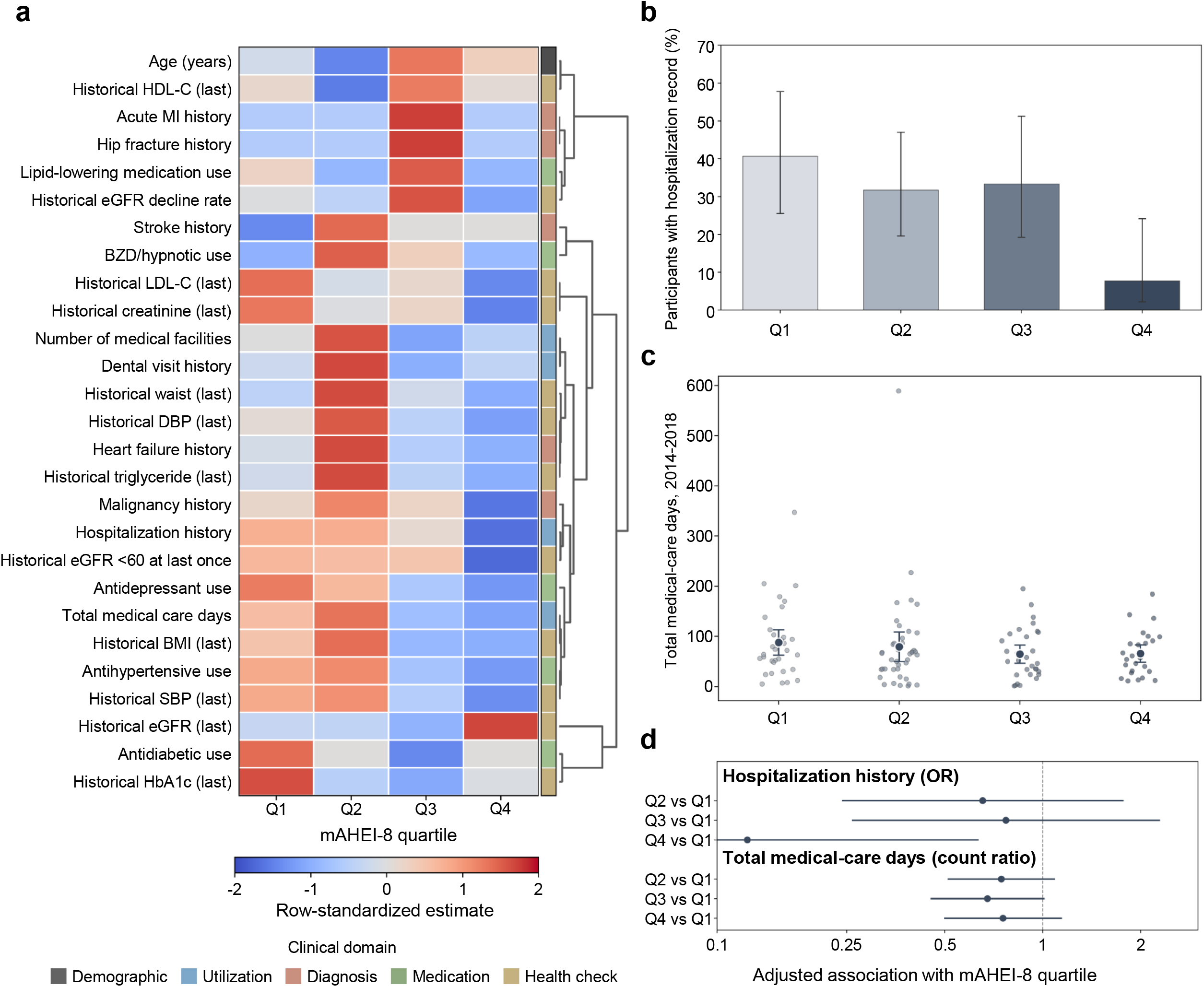
Higher mAHEI-8 corresponds to a more favorable preceding health profile. Preceding health-insurance claims and municipal health-check information were obtained during 2014–2018; participant age is included in panel a as a demographic reference. a, Claims-derived and municipal health-check profiles among participants with mAHEI-8. Each cell represents a quartile-specific proportion or mean standardized within that variable. For claims-derived event and healthcare-utilization variables, participants with no identified medical claim during 2014–2018 were assigned zero under this descriptive specification, whereas missing municipal health-check measurements were not imputed. Rows were hierarchically clustered using correlation distance and average linkage, with Q1–Q4 kept in fixed column order. b–d, Healthcare utilization among 129 participants with at least one identified medical claim, analyzed without zero filling. b, Proportion with at least one hospitalization during 2014–2018, with Wilson 95% confidence intervals (Q1–Q4: 13/32, 13/41, 10/30 and 2/26). c, Total medical-care days; small points represent individual participants, and large points and error bars indicate means and 95% confidence intervals. d, Age- and sex-adjusted associations of Q2, Q3 and Q4 with hospitalization and total medical-care days, with Q1 as the reference category. Hospitalization was analyzed using logistic regression and is shown as odds ratios; total medical-care days were analyzed using negative-binomial regression (NB2) and are shown as count ratios. Points indicate estimates and horizontal lines indicate 95% confidence intervals.

Among the 129 participants with at least one identified medical claim, the proportion with at least one hospitalization during 2014–2018 was 40.6%, 31.7%, 33.3% and 7.7% across Q1–Q4, respectively, and was lowest in Q4 (Fig. 3b). Mean total medical-care days during the same period were 87.7, 79.1, 64.5 and 65.6 days (medians, 68, 65, 51 and 59 days, respectively), tending to be lower in the higher-mAHEI-8 groups (Fig. 3c). However, the distribution was strongly right-skewed, and between-quartile differences were small relative to the extent of inter-individual variation. After adjustment for age and sex, the odds ratios for hospitalization relative to Q1 were 0.65, 0.77 and 0.12 for Q2, Q3 and Q4, respectively, whereas the corresponding count ratios for total medical-care days were 0.75, 0.68 and 0.76 (Fig. 3d). The clearest adjusted association was observed for hospitalization in Q4, whereas estimates for total medical-care days were lower across Q2–Q4 but remained statistically uncertain. Because these health records preceded the FY2022 dietary assessment, these associations do not establish causal direction, but indicate that the currently observed mAHEI-8 gradient also corresponded to healthcare utilization and clinical status several years earlier.

### mAHEI-8 did not define a discrete contemporary multivariate phenotype

Given the correspondence of mAHEI-8 with healthcare utilization and health-check measures from 2014–2018, we next examined how these differences in diet quality were represented in the contemporary biological state. Specifically, using blood and urine tests, serum metabolomics and Raman data acquired contemporaneously in FY2022, we assessed the correspondence between mAHEI-8 and each multivariate phenotype. In the blood/urine, serum metabolome, Raman and integrated PHATE spaces, mAHEI-8 quartiles broadly overlapped and showed no clear separation (Fig. 4a). The same pattern was observed with PCA and UMAP visualizations (Supplementary Fig. 4).

**Fig. 4.**
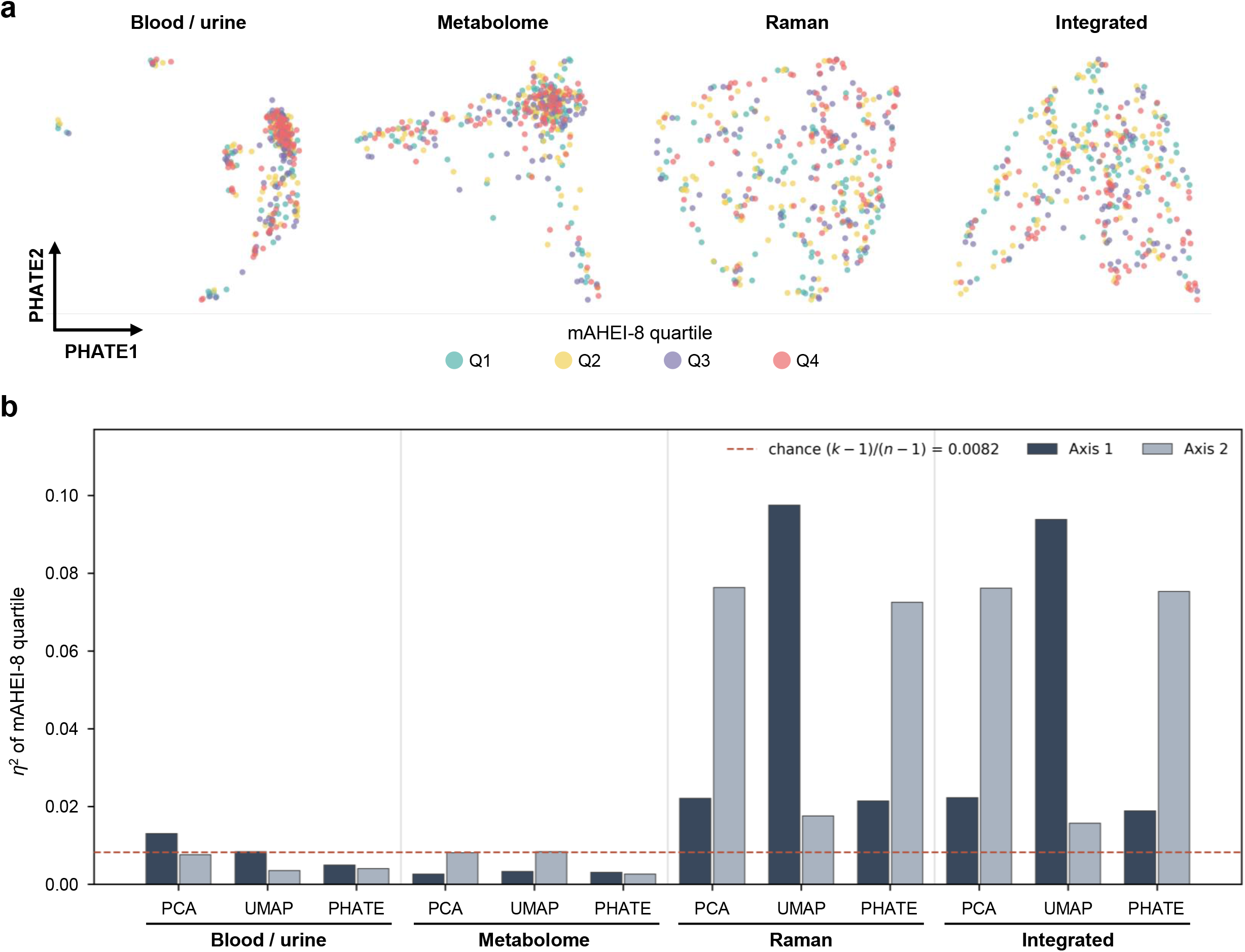
mAHEI-8 quartiles do not form discrete groups in contemporary multivariate phenotypes. The four phenotypic representations were constructed among 370 participants with complete blood/urine, serum metabolome and quality-controlled Raman data. a, PHATE representations coloured by mAHEI-8 quartile for the 367 participants with corresponding dietary data. Dietary variables were not used to construct the embeddings. Corresponding PCA and UMAP representations are shown in Supplementary Fig. 4. b, η² for mAHEI-8 quartile on each axis across the four phenotypic spaces and three dimensionality-reduction methods. The dashed line indicates the expected value under random labelling, (k − 1)/(n − 1) = 0.0082. η² is used as a descriptive statistic; permutation-based P values were not calculated.

When the correspondence between each low-dimensional axis and mAHEI-8 quartile was quantified using η^2^, values for the blood/urine and metabolome axes were generally close to the chance-level reference, whereas some axes in the Raman and integrated phenotype spaces showed relatively larger values (Fig. 4b). Thus, although mAHEI-8 was not a major source of variation separating multilayer phenotypes into clearly discrete states, some diet-related variation embedded within the composite score may have been represented as continuous variation, particularly in phenotype spaces that included Raman information. This raised the possibility that more specific dietary information could be present in the Raman phenotype even when the composite score itself did not define a clear state.

### Serum Raman added complementary information for specific dietary variables

To test whether such specific dietary information was represented as continuous variation, particularly in Raman-containing phenotype spaces, we evaluated 28 variables drawn from BDHQ-derived foods and nutrients, the total mAHEI-8 score and its components. These 28 variables were prespecified before the cross-modal screen and were all reported irrespective of the results. Using a common cross-validation procedure, we quantified the incremental predictive performance obtained by adding Raman to a model based on blood tests and serum metabolomics (ΔOOF R²). Improved predictive performance after adding Raman remained supported after FDR correction for 17 of the 28 variables (Fig. 5a). ΔOOF R² was 0.139 for coffee, 0.059 for β-carotene equivalent, 0.057 for green tea, 0.054 for total mAHEI-8, 0.044 for vegetables, 0.037 for dietary fiber and 0.036 for other vegetables, indicating that the additional predictive information provided by Raman was not confined to a single food or nutrient variable. In negative controls generated by randomly shuffling the outcomes, ΔOOF R² remained close to zero (Supplementary Fig. 5). In a sensitivity analysis that added age and sex to the baseline model, the Raman increment remained supported after FDR correction for 14 of the 17 variables that met the FDR criterion in the primary analysis. Overall, 15 variables met the FDR criterion in the sensitivity analysis, including β-cryptoxanthin, which had not met the criterion in the primary analysis. For β-carotene equivalent, ΔOOF R² changed only slightly, from 0.059 in the primary analysis to 0.053 after the addition of age and sex to the baseline model (q = 0.0028; Fig. 5a). Thus, most of the additional predictive information provided by Raman was not explained solely by variation associated with age and sex.

**Fig. 5.**
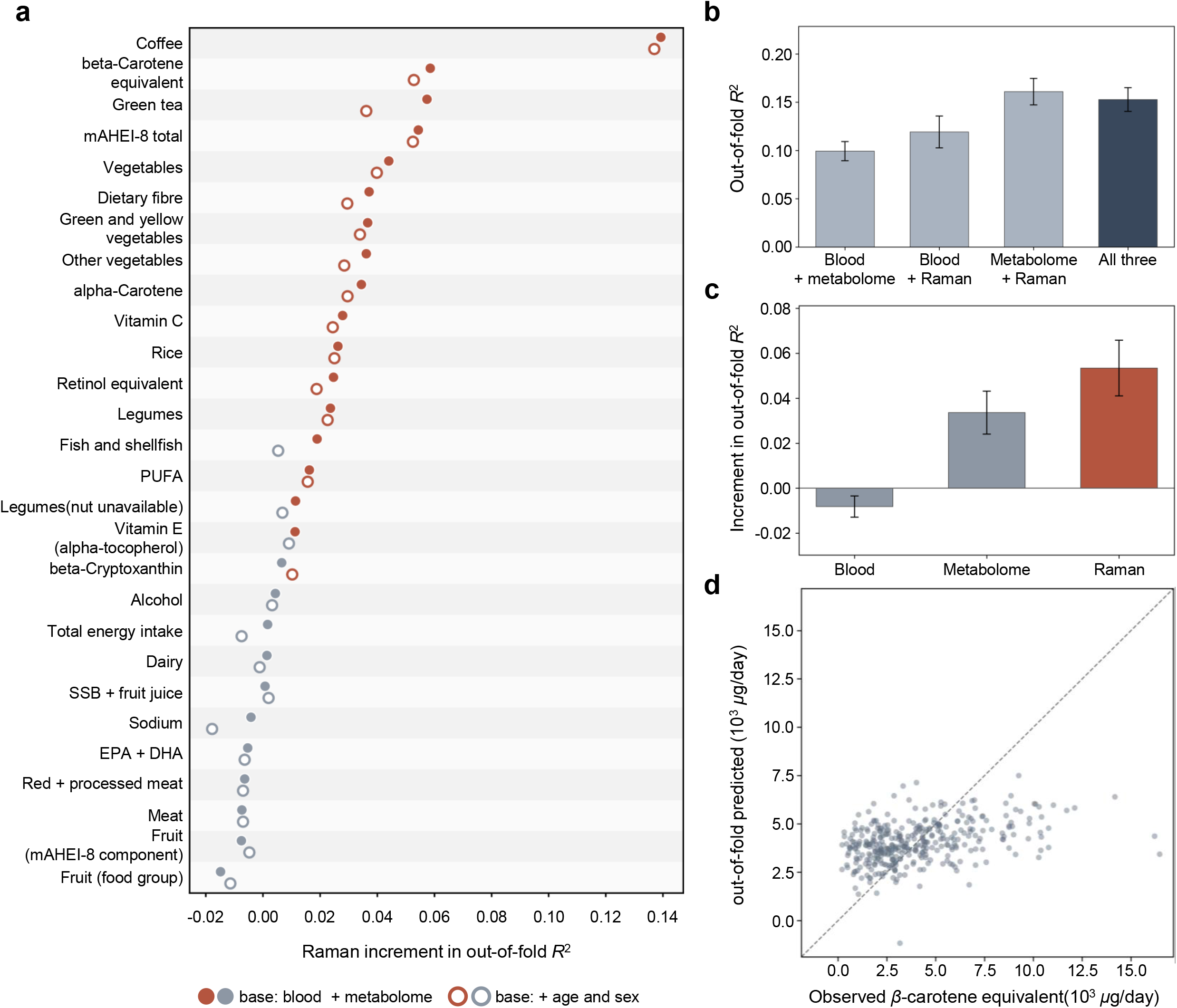
Serum Raman provides complementary predictive information for multiple dietary variables. All models were evaluated in the common analytical population of 367 participants. a, ΔOOF R² obtained by adding Raman to the blood + metabolome model for 28 dietary variables prespecified before the cross-modal screen and retained for reporting. Urine measurements were not included in these predictive models. Outcomes are ordered by ΔOOF R² in the primary analysis. Filled and open circles represent the primary analysis and the age- and sex-adjusted sensitivity analysis, respectively. P values were obtained from null distributions generated by 2,000 permutations of the Raman block and were corrected across the 28 outcomes using the Benjamini–Hochberg procedure. Red circles indicate variables with a Benjamini–Hochberg-adjusted permutation P value (q < 0.05), and grey circles indicate variables that did not meet this criterion; color is assigned separately for the primary and age- and sex-adjusted analyses. Negative controls are shown in Supplementary Fig. 5. b, OOF R² for four modality combinations predicting β-carotene equivalent. c, Incremental OOF R² after removing each layer from the three-layer model. Error bars in b,c show the s.d. across 10 cross-validation repeats and are not confidence intervals. d, OOF-predicted versus BDHQ-derived β-carotene equivalent from the three-layer model. Each point represents the prediction for a held-out participant.

Among the variables supported in this screen, β-carotene equivalent was of particular interest because the Raman information associated with it could be traced to established molecular vibrations characteristic of carotenoids^17^ and was therefore examined in greater detail. In repeated out-of-fold ridge regression among the common set of 367 participants, the OOF R² for β-carotene equivalent was 0.099 using blood tests and serum metabolomics and 0.153 using all three phenotype layers (difference in OOF R² = 0.053, repeat SD = 0.012; Fig. 5b). When each phenotype layer was removed in turn from the three-layer model combining blood tests, serum metabolomics and Raman, OOF R² decreased by 0.053 after removal of Raman and by 0.034 after removal of serum metabolomics. By contrast, the model without blood tests performed slightly better than the three-layer model (difference in OOF R^2^ = −0.008; Fig. 5c). Raman and serum metabolomics therefore provided predictive information for β-carotene equivalent that was not completely overlapping. The Spearman rank correlation between out-of-fold predicted and observed values from the three-layer model was ρ = 0.381 (P = 3.8 × 10^−14^; Fig. 5d). Predictions tended to underestimate β-carotene equivalent at higher intake levels, but adding visit date had little effect on out-of-fold predictive performance (Supplementary Fig. 6a–d). The incremental predictive performance of Raman was also retained after applying a log(1+x) transformation to the outcome to reduce the influence of high values (Supplementary Fig. 6e).

### Carotenoid-associated Raman variation reflected a broader plant-food-related dietary dimension

Having established that Raman provided complementary predictive information for β-carotene equivalent, we next examined its spectroscopic basis. Carotenoids exhibit strong Raman scattering arising from their conjugated double-bond systems, with characteristic bands near 1004–1006, 1158 and 1518 cm^−117^. The band near 1158 cm^−1^ is attributed primarily to C–C stretching, whereas the band near 1518 cm⁻¹ corresponds to C=C stretching. Because the 1004–1006 cm⁻¹ region may also contain contributions from the phenylalanine ring-breathing vibration in serum^18^, we restricted molecular interpretation in this study to the regions near 1158 and 1518 cm^−1^.

Mean Raman spectra across mAHEI-8 quartiles were globally similar, but quartile-related variation was evident around major Raman bands, including those near 1158 and 1518 cm⁻¹ (Fig. 6a). We selected the spectral variables nearest to 1158 and 1518 cm⁻¹ on the common Raman-shift grid and defined their equal-weight average as a two-band score. This score was positively rank-correlated with BDHQ-derived β-carotene equivalent (Spearman ρ = 0.185, 95% CI 0.084–0.282, P = 3.5 × 10^−4^, n = 370; Fig. 6b). The individual bands showed associations in the same direction, with ρ = 0.189 at 1158 cm^−4^ and ρ = 0.181 at 1518 cm^−1^ (both P < 5 × 10^−4^; Fig. 6c). These associations were retained after adjustment for age and sex (Fig. 6c).

**Fig. 6.**
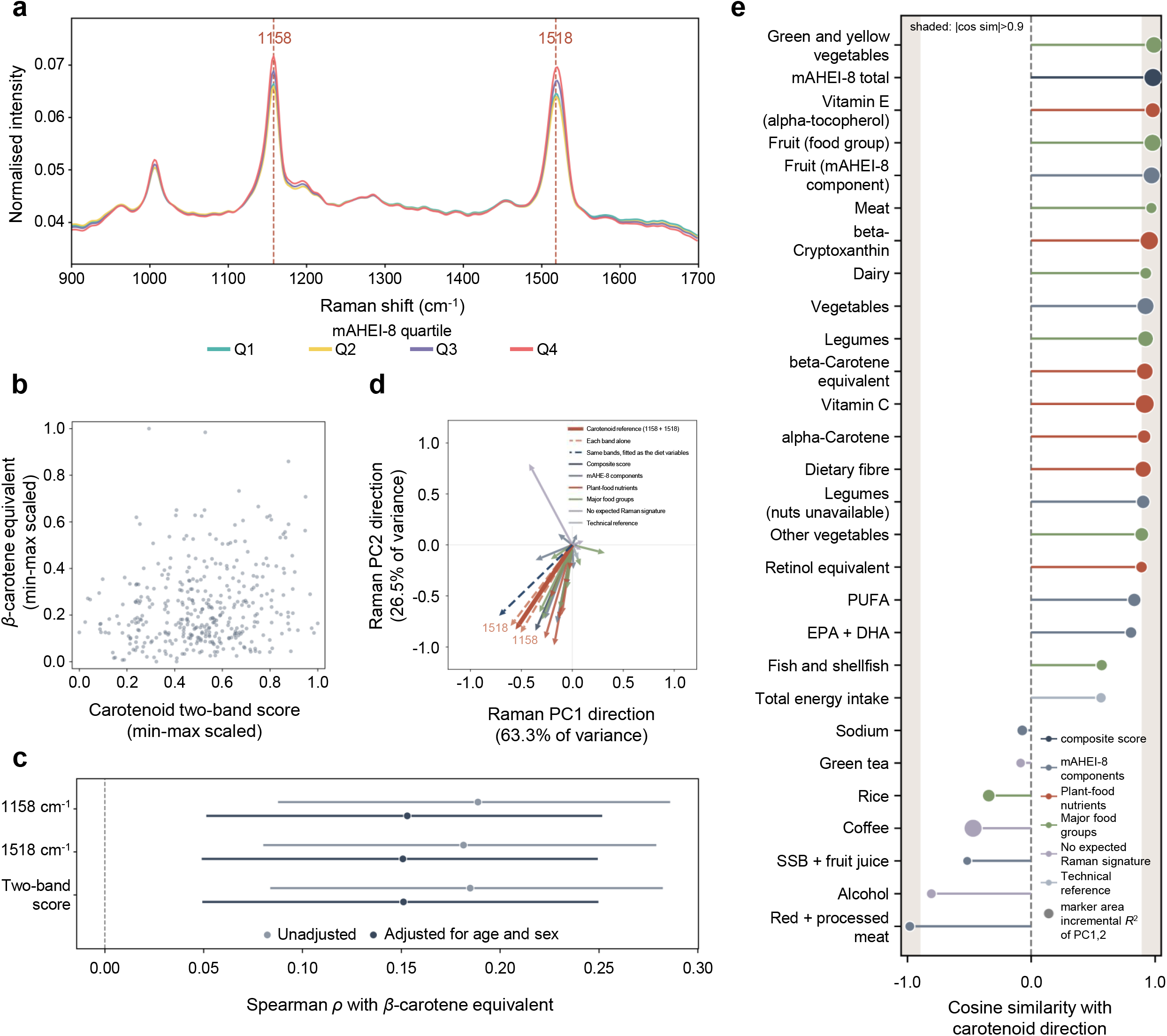
Carotenoid-assigned Raman variation is shared with a broader plant-food-related dietary dimension. a–c, Data from 370 participants with quality-controlled Raman spectra and corresponding dietary information. a, Mean serum Raman spectra by mAHEI-8 quartile. Dashed lines indicate the prespecified 1158 and 1518 cm⁻¹ regions. b, Equal-weight two-band score of the 1158/1518 cm⁻¹ intensities versus BDHQ-derived β-carotene equivalent. c, Spearman correlations for each band and the two-band score, shown as unadjusted and age- and sex-adjusted estimates. d,e, Directions in the Raman PC1–PC2 plane. Raman PCA and the loading-based carotenoid reference direction were derived from all 373 quality-controlled Raman spectra; dietary association directions were estimated in participants with corresponding dietary and demographic data (typically n = 370). The carotenoid reference is the unit vector defined from PCA loadings at 1158 and 1518 cm⁻¹. Dietary directions are age- and sex-adjusted association directions in the same plane (standardized dietary variable ∼ PC1 + PC2 + age + sex); arrow length and marker size are proportional to the incremental R² explained by PC1 and PC2 beyond age and sex. e, Cosine similarity of each dietary direction with the loading-based carotenoid reference direction. These directional analyses are descriptive; inferential P values were not calculated.

To confirm that focusing on these two established bands did not simply reflect post hoc selection of regions with the strongest correlations, we also surveyed diet-related structure across the full spectrum. Separately from the 28 variables used in the cross-modal screen, we constructed a descriptive correlation map for all pairwise combinations of 70 available BDHQ food variables and 549 Raman shifts. Diet-related correlations were broadly distributed across the spectrum, and 1158 and 1518 cm^−1^ were not empirically the regions with the largest correlations (Supplementary Fig. 7). Thus, the correspondence between β-carotene equivalent and these two regions was not a consequence of retrospectively selecting the most strongly correlated spectral regions, but was consistent with established molecular assignments for carotenoids.

We further asked whether this molecular correspondence was specific to β-carotene equivalent or shared with broader diet-related variation. In Raman PCA, PC1 and PC2 together explained 89.8% of the total variance (Fig. 6d; PC1, 63.3%; PC2, 26.5%). The carotenoid reference direction defined from the PCA loadings at 1158 and 1518 cm⁻¹ closely matched the direction obtained when the two-band score was entered into the same model used for the dietary variables (cosine similarity = 0.978). When the 28 diet-related variables from the cross-modal screen were positioned in the PC1–PC2 plane, many plant-food- and nutrient-related variables, together with total mAHEI-8 and its vegetable, fruit and legume components, aligned with the carotenoid reference direction (Fig. 6d). By contrast, red + processed meat, alcohol, sugar-sweetened beverages + fruit juice, rice and coffee were positioned in the opposite direction (Fig. 6d). The incremental R^2^ explained by PC1 and PC2 beyond age and sex was relatively large for vitamin C (0.085), β-cryptoxanthin (0.081), total mAHEI-8 (0.078), coffee (0.077), β-carotene equivalent (0.063) and green and yellow vegetables (0.063). Although coffee showed substantial variation along PC1 and PC2, its association direction was negatively aligned with the carotenoid reference direction (cosine similarity = −0.469), in contrast to the positive alignment observed for several plant-food- and carotenoid-related variables (Fig. 6e). Thus, the Raman variation corresponding to β-carotene equivalent was shared with a broader plant-food-related dietary dimension that included mAHEI-8, whereas coffee-related Raman variation represented a distinct direction within the same Raman space.

## Discussion

In this study, we linked habitual dietary assessment at the individual level to temporally preceding health information and to multiple contemporaneous molecular and physiological phenotypes to examine how overall diet quality and more specific dietary information are represented in biological phenotypes. The mAHEI-8 gradient was driven unevenly by a subset of its components, and higher mAHEI-8 corresponded to a more favorable preceding health background among participants for whom linkage was possible. Although mAHEI-8 quartiles did not separate contemporary multidimensional phenotypes into clearly discrete states, some low-dimensional axes showed continuous variation corresponding to diet quality, particularly in phenotype spaces involving Raman. Extending the analysis to specific food and nutrient variables showed that Raman provided additional predictive information beyond blood tests and serum metabolomics for multiple variables, and for β-carotene equivalent, this information could be traced from established carotenoid-assigned bands to a broader plant-food-related dietary dimension.

The mAHEI-8 used here is a study-specific adaptation of AHEI-2010 and is not directly interchangeable with the original score. Nevertheless, previous studies have used reduced versions of AHEI-2010 according to the dietary information available and the intake characteristics of the target population. An eight-component modified AHEI-2010 was used in a large prospective cohort in Shanghai^19^, and the JACC Study in Japan used a 10-component modified AHEI-2010 because whole-grain intake was unavailable^3^. A study comparing several diet-quality indices, including AHEI-2010, in Japanese adults also found that AHEI-2010 was associated with a lower prevalence of inadequate intake of multiple nutrients^20^. In the present population, approximately 82% of the Q1–Q4 score difference arose from four components, and EPA + DHA showed a pronounced ceiling effect. It is therefore important to interpret mAHEI-8 not as a uniform dietary exposure, but together with the specific combination of foods and nutrients that generated its gradient. The correspondence of mAHEI-8 with historical hospitalization and several cardiometabolic and renal measures provides health context for the dietary differences captured by the score; however, because the health information was from 2014–2018 whereas dietary assessment was conducted in FY2022, this correspondence should be interpreted as an association with longer-term health background rather than as a causal relationship.

Molecular signatures associated with healthy dietary patterns have been reported previously. In a controlled-feeding study, urinary metabolic phenotypes distinguished between different dietary patterns^21^, and in free-living populations, urinary metabolomics has been used to predict dietary patterns with replication in an independent population^22^. Circulating metabolic signatures associated with AHEI-2010 have also been identified in multiple cohorts^8^K^9^, and plasma proteomic signatures corresponding to healthy dietary patterns have been reported^23^. These prior findings are compatible with our observation that mAHEI-8 quartiles did not separate clearly in PCA, UMAP or PHATE spaces. Many previous studies used supervised approaches to identify or predict molecular features associated with diet, whereas our unsupervised analyses asked whether overall diet quality formed a major axis of variation in the phenotype as a whole. If dietary effects are distributed weakly across many features, and if the same mAHEI-8 score can arise from different combinations of components, reproducible diet-associated signatures may exist without quartiles forming natural clusters. Indeed, despite weak global separation of quartiles in the present study, adding Raman improved prediction for multiple dietary variables, including mAHEI-8 itself. These results suggest that diet-related biological information may be represented as continuous variation distributed across multiple phenotypic layers rather than as discrete biological states.

There is also precedent for detecting nutrition-related information using Raman spectroscopy. Carotenoid measurements by skin resonance Raman spectroscopy have been validated against circulating and skin carotenoids and fruit and vegetable intake^10^K^11^K^24^, and associations with diet quality based on the Healthy Eating Index have also been reported^25^. Changes in human serum Raman spectra after carotenoid supplementation have likewise been observed^26^. The novelty of the present study therefore does not lie in showing that Raman spectroscopy is related to carotenoids or diet per se. Rather, we show in free-living individuals consuming their habitual diets that serum Raman spectra provide additional predictive information beyond blood tests and serum metabolomics for multiple dietary variables, and that information corresponding to β-carotene equivalent can be traced to established molecular vibrations. The correlation between the two-band score and BDHQ-derived β-carotene equivalent was modest, but habitual intake estimated from questionnaires is not expected to correspond one-to-one with circulating biomarkers. In the JPHC validation study, correlations between questionnaire-derived β-carotene intake and serum β-carotene concentration were 0.28 in men and 0.11 in women^27^. Because individual serum carotenoids were not directly quantified in the same samples in our study, the 1158/1518 cm^−1^ features should be interpreted as carotenoid-assigned bands rather than as direct quantification of circulating carotenoid concentrations.

Coffee showed the largest increment in predictive performance after Raman was added to the blood and metabolome model. Coffee was included in the fixed dietary-variable panel before the cross-modal screen, and its result was reported irrespective of its magnitude. However, unlike β-carotene equivalent and several plant-food-related variables, the association direction for coffee in Raman PC1–PC2 space was negatively aligned with the prespecified carotenoid reference direction. Thus, the strong Raman-related information for coffee did not simply reflect the carotenoid-associated variation identified for plant-food-related variables. Because no a priori spectroscopic assignment was specified for coffee, and coffee intake may be associated with multiple chemical, metabolic and behavioural factors, we did not pursue a molecular interpretation of this association.

The carotenoid-associated Raman variation was also not confined to β-carotene equivalent. Along the reference direction defined from 1158/1518 cm^−1^, several plant-food- and nutrient-related variables aligned, including β-carotene equivalent, vitamin C, β-cryptoxanthin, green and yellow vegetables and total mAHEI-8. Accordingly, serum Raman is better viewed not as a direct readout of a single nutrient or of a ‘healthy diet’ as a whole, but as a phenotypic layer that captures particular molecular and food-related dimensions embedded within complex dietary exposure. This perspective shifts the analytical focus from seeking overall diet quality as a single biological state to decomposing the information contained within a summary score into biologically traceable dimensions and matching those dimensions to complementary phenotypic layers. The present study does not comprehensively map the full set of such dimensions, but it indicates that moving from composite diet scores towards biologically traceable dietary dimensions may provide a useful framework for understanding how diet corresponds to an individual’s biological state.

This study has several limitations. There was a temporal gap between the preceding health information and the FY2022 dietary assessment, precluding inference about causal direction from their correspondence. Participants with municipal records identified during 2014–2018 were older than the overall population, limiting the generalizability of analyses based on the subset with preceding health information, particularly municipal health-check measurements and medical claims. Analyses of contemporaneous diet and molecular phenotypes were also observational and cross-sectional, and BDHQ-based dietary assessment is subject to measurement error. mAHEI-8 is a study-specific modification and cannot be directly compared with the original AHEI-2010. In addition, individual serum carotenoids were not directly quantified in the same samples, and Raman spectra may be influenced by technical factors including measurement date and run-dependent Raman-shift axes. Although we applied common-axis correction, technical quality control and sensitivity analyses, independent validation across different populations and measurement settings will be necessary to establish the generalizability of the Raman–diet correspondence.

In conclusion, diet quality represented by mAHEI-8 corresponded to a more favorable preceding health background, whereas its quartiles did not separate contemporary multidimensional phenotypes into a single clear state. More specific diet-related information was nevertheless detectable in biological phenotypes, and serum Raman provided plant-food-related information complementary to other measurement layers. Part of this information could be traced, through its correspondence with β-carotene equivalent, to established carotenoid-assigned bands and was further shared with a broader plant-food-related dietary dimension. Together, these findings suggest that habitual diet is represented not as a single biological state, but through biologically traceable dietary dimensions distributed across complementary phenotypic layers.

## Methods

### Study design and participants

This observational study used data from the Tsukuba Happiness Life Study^12−14^, which includes community-dwelling middle-aged and older adults in Japan. The primary analyses used dietary assessment, anthropometric and clinical measurements, blood and urine tests, serum metabolomics and serum Raman spectra obtained in the FY2022 survey. To assess preceding health background, medical-insurance claims and municipal health-check records held by Tsukuba City from 2014–2018 were linked at the participant level.

Among the 405 participants who underwent the FY2022 health examination, BDHQ data were available for 401, for whom mAHEI-8 could be calculated. After Raman quality control (QC), 373 spectra remained. Fig. 4 embeddings were constructed for 370 participants with complete blood/urine, metabolome and QC Raman data; mAHEI-8 overlays and η^2^ analyses used the 367 participants with corresponding dietary data. Fig. 5 used the same 367 participants. In Fig. 6, Raman PCA and the loading-based carotenoid reference used all 373 QC spectra, whereas band–diet associations and dietary directions used participants with corresponding dietary and demographic data (typically n = 370).

### Ethics

The study was approved by the ethics committee of the University of Tsukuba Hospital (approval no. R03-335; June 21, 2022). The study was conducted in accordance with the Declaration of Helsinki and relevant institutional guidelines, and written informed consent was obtained from all participants.

### Dietary assessment and construction of mAHEI-8

Habitual dietary intake was assessed using the BDHQ. Food-group and nutrient intakes and carotenoid-related variables were derived from the FY2022 BDHQ output. β-Carotene equivalent was calculated as β-carotene + 0.5 × α-carotene + 0.5 × β-cryptoxanthin.

Overall diet quality was assessed using an 8-component modified AHEI-2010 score (mAHEI-8; range 0–80) based on AHEI-2010. The components were vegetables, fruit, legumes, EPA + DHA, polyunsaturated fatty acids (PUFA), sugar-sweetened beverages + fruit juice, red + processed meat and sodium, each scored from 0 to 10. The principal modifications from the original AHEI-2010 were that nuts + legumes were represented by legumes because nuts could not be independently quantified using the BDHQ; sugar-sweetened beverages + fruit juice was calculated from cola and 100% fruit juice; and whole grains, trans fat and alcohol were excluded. Sodium was scored according to sex-specific intake deciles. mAHEI-8 is a study-specific adaptation and was not considered directly interchangeable with the original AHEI-2010.

mAHEI-8 was treated as a continuous variable where appropriate, whereas quartiles were used for descriptive analyses and visualization. In the dietary-direction analysis in the Raman PCA plane, BDHQ variables with fewer than 30 non-missing observations or zero variance were excluded.

### Preceding health-insurance claims and municipal health-check records

From medical-insurance claims during 2014–2018, hospitalization history, total medical-care days, disease history, medication use, medical procedures, number of therapeutic drug classes, number of healthcare institutions visited and other healthcare-utilization variables were derived at the participant level and aggregated across the observation period. Where distinguishable in the source data, suspected and confirmed diagnoses were treated separately.

From municipal health-check records, we obtained BMI, waist circumference, systolic and diastolic blood pressure, HbA1c, LDL cholesterol, HDL cholesterol, triglycerides, serum creatinine and eGFR. For variables designated as ‘last’, the final available measurement during the observation period was used.

For the descriptive heatmap in Fig. 3a, binary variables were summarized as proportions and continuous variables as means within each mAHEI-8 quartile. For claims-derived event and healthcare-utilization variables, participants with no identified medical claim during 2014–2018 were assigned zero under this descriptive specification, whereas missing municipal health-check measurements were not imputed. Quartile estimates were then standardized within each variable. Rows with estimates available across all four quartiles were hierarchically clustered using correlation distance and average linkage, while the Q1–Q4 column order was fixed.

For Fig. 3b–d, healthcare-utilization analyses were restricted to the 129 participants with at least one identified medical claim and were performed without zero filling. For the adjusted analyses in Fig. 3d, hospitalization history was analyzed using logistic regression and total medical-care days using negative-binomial regression (NB2), with mAHEI-8 quartile entered as a categorical variable with Q1 as the reference and adjustment for age and sex. Because the Poisson model showed marked overdispersion, the negative-binomial model was used for total medical-care days.

### Routine blood and urine tests

Routine blood and urine measurements from FY2022 were used as the clinical phenotype layer. Visit date was defined as the date of the FY2022 health examination, corresponding to the blood collection date, and was distinct from the Raman measurement date. Participant identifiers, visit date, age and sex were excluded from the phenotypic features. Numerical features with ≥20% missingness or zero variance were removed, and remaining missing values were imputed with the median. For qualitative urine tests with <20% missingness, the five levels (−), (±), (1+), (2+) and (3+) were one-hot encoded after missing values were imputed with the mode. Both blood and urine measurements were used for the multivariate phenotype representations, whereas only blood measurements were used in the cross-modal prediction analyses described below.

### Serum metabolomics

Serum metabolomic profiling was performed using serum collected in FY2022. Metabolites were extracted from 50 μl of each serum sample, derivatized by methoximation and trimethylsilylation, and analyzed by gas chromatography–mass spectrometry (GC–MS; GCMS-TQ8030, Shimadzu, Japan). The peak area of each metabolite divided by the peak area of the internal standard, 2-isopropylmalic acid, was used as the analytical feature. These values were treated as relative amounts within each metabolite and were not used to compare absolute abundance across metabolites.

Quality-control (QC) samples were measured at regular intervals throughout the analytical sequence. Features with a QC coefficient of variation ≥25% or meeting analytical exclusion criteria were removed, leaving 106 QC-passed features for analysis. Values were converted to numeric form, verified to be non-negative and transformed using log(1+x) before downstream analysis.

### Serum Raman spectroscopy, preprocessing and quality control

Serum samples stored at −80 °C were thawed on ice for 30 min and gently mixed. After brief centrifugation, 80 μl was dispensed into an optical-grade quartz-bottom 96-well plate. Samples were equilibrated for 10 min at room temperature (22–25 °C) before measurement.

Raman spectra were acquired using a custom Raman microscope comprising an STR-Raman imaging module (AIRIX Corp., Japan) integrated with an inverted microscope (Ti2-E, Nikon, Japan). A 532-nm continuous-wave laser (MLL-U-532A-100mW, CNI, China) was delivered through a 20× objective lens (NA 0.50; working distance 2.1 mm), with a laser power of 20 mW at the sample surface. Backscattered Raman signals were transmitted through a 50-μm-core optical fiber to a spectrometer equipped with a 1200 grooves mm⁻¹ grating (Kymera 193i, Andor-Oxford Instruments, UK) and detected with a CCD detector cooled to −60 °C (iVac 316-LDC-DD, Andor-Oxford Instruments, UK) using a 120-s exposure. Two replicate spectra were acquired for each participant, and the Raman-shift axis was calibrated before each measurement session using the 884, 1455 and 2927 cm⁻¹ ethanol bands.

Cosmic-ray artefacts were removed using AIRIX software. To correct differences in the Raman-shift grid among acquired spectra, a common grid was constructed from the individual axes and the 600–1700 cm⁻¹ region was mapped at 2 cm⁻¹ intervals. Each replicate underwent Savitzky–Golay smoothing (window length 9, third-order polynomial, derivative 0)²⁸ followed by L2 normalization. The two replicates were then averaged with equal weight, and the participant-level mean spectrum was L2-normalized again. This axis-correction–Savitzky–Golay–L2 procedure was used as the primary preprocessing pipeline for downstream analyses.

Technical quality control assessed peak-top flatness around carotenoid-associated bands across all 376 spectra. Fifty spectra were flagged as showing relatively pronounced peak-top flatness in the technical QC assessment, whereas three spectra with severe plateauing were excluded before construction of the primary Raman matrix. The technical characteristics of peak-top flatness were additionally evaluated across all 376 spectra (Supplementary Fig. 8). Flatness classification was sensitive to the Raman-shift grid and raw/background intensity and was largely resolved after primary preprocessing. No spectra beyond the three excluded by technical quality control were additionally removed on the basis of flatness. Technical sensitivity analyses using alternative preprocessing schemes were also performed, and the preprocessing method was not selected on the basis of the strength of association with diet.

### Multivariate phenotypic representations

For each feature matrix—blood/urine, serum metabolome and Raman—participant identifiers, visit date, age and sex were excluded, and feature-wise standardization was applied after handling missing values. Principal-component analysis (PCA)²⁹ was performed and the minimum number of principal components required to reach 90% cumulative explained variance was retained. PHATE³⁰ was applied to the retained PC scores with n_components = 2, n_pca = None, knn = 5, t = ‘auto’, decay = 40 and random_state = 0. UMAP³¹ was applied with n_neighbors = 15, min_dist = 0.1, Euclidean distance and random_state = 0. Dietary variables were not used to construct these representations; mAHEI-8 quartiles were overlaid as external labels after embedding construction.

For the integrated representation, participant-aligned blood/urine, metabolome and Raman blocks were preprocessed and standardized within each block, concatenated without block-specific weighting, and analyzed by PHATE after global PCA. The correspondence between each embedding axis and mAHEI-8 quartile was summarized using one-way η². For four groups and n = 367, the expected value under random labelling was (k − 1)/(n − 1) = 0.0082, which was used as a descriptive reference in Fig. 4b. Permutation-based P values were not calculated.

### Cross-modal prediction of dietary variables

To assess whether Raman contained diet-related information beyond routine blood tests and serum metabolomics, we analyzed a fixed panel of 28 dietary variables. Unlike the multivariate phenotype representations, which used the combined blood/urine clinical phenotype layer, the predictive baseline models used blood measurements only; urine measurements were not included. Accordingly, ‘blood’ in the prediction models refers to routine blood measurements only.

The panel was defined before the cross-modal screen to cover complementary aspects of dietary exposure rather than to achieve a target number of outcomes. It comprised total mAHEI-8 and all eight components, selected plant-food-related nutrients and major food groups, green tea intake from the BDHQ food variable BEV1, coffee intake from BEV3, alcohol intake from the BDHQ nutrient variable ALC, and total energy intake as a reference for overall intake. All variables meeting the prespecified availability and variance criteria were retained and reported irrespective of the screening results.

For the cross-modal screen, ridge models were evaluated using shuffled 5-fold outer cross-validation repeated 10 times. Within each outer training fold, the ridge penalty for the baseline model was selected by inner 5-fold cross-validation. Feature filtering, missing-value imputation and standardization were re-estimated separately within each inner training fold and applied to the corresponding inner validation fold. After penalty selection, preprocessing and model fitting were repeated using the complete outer training fold, and predictions were generated for the held-out outer fold. The same outer-fold partition and fold-specific ridge penalty selected for the baseline model were then used for the corresponding Raman-augmented model and all Raman-block permutation fits, with preprocessing re-estimated from each outer training fold. Out-of-fold predictions were averaged across the 10 outer-cross-validation repeats before R² and ΔOOF R² were calculated.

The primary statistic was ΔOOF R² = R²(blood + metabolome + Raman) − R²(blood + metabolome). For each outcome, a null distribution of ΔOOF R² was constructed from 2,000 permutations in which the participant rows of the Raman block were shuffled while the outcome and the blood and metabolome blocks were kept fixed. The predictive model was refitted for each permutation using the same outer-fold partition and fold-specific ridge penalty as in the observed analysis. One-sided permutation P values were calculated as (1 + k)/(1 + B), where k was the number of permuted ΔOOF R² values at least as large as the observed value and B = 2,000. Benjamini–Hochberg false-discovery-rate correction was applied across the 28 outcomes within each analysis³². An analysis in which age and sex were included as additional covariates in both the baseline and Raman-augmented models was treated as a post hoc sensitivity analysis. Ten shuffled outcomes were analyzed using the same cross-validation and Raman-block permutation procedures as negative controls.

### Multimodal prediction of β-carotene equivalent

For β-carotene equivalent, four modality combinations—blood + metabolome + Raman, blood + metabolome, blood + Raman and metabolome + Raman—were compared in the common cohort of 367 participants. As defined above, the blood block comprised routine blood measurements and did not include urine measurements. All models used ridge regression³³. Ten shuffled 5-fold outer-cross-validation partitions were generated using different random seeds, and the same set of partitions was applied to all modality combinations. Feature selection, missing-value imputation and standardization were all estimated within the training fold. Features were retained if they were non-missing in at least 80% of training observations and had finite non-zero variance; remaining missing values were imputed using the training median. For this descriptive comparison, the ridge penalty α was selected separately for each modality combination from 41 candidates spanning 10⁻⁴ to 10⁴ using inner 5-fold cross-validation. Predictions across the ten outer-cross-validation repeats were averaged for each participant.

The primary performance metric was OOF R², with RMSE, MAE and observed–predicted Spearman correlation as secondary metrics. The change associated with each phenotypic layer was quantified as the difference in OOF R² between the three-layer model and the two-layer model excluding that layer, such that a positive value indicated lower performance after removal of the layer. Model diagnostics were based on the averaged OOF predictions obtained from the repeated nested cross-validation. Sensitivity models that additionally included visit date or used log(1+x)-transformed β-carotene equivalent as the outcome were fitted using the same model-specific nested cross-validation procedure.

### Carotenoid-associated Raman analyses

The spectral variables nearest to 1158 and 1518 cm⁻¹ on the corrected Raman grid were selected a priori on the basis of established carotenoid Raman assignments¹⁷K²⁶. The 1004–1006 cm⁻¹ region was excluded from the primary molecular interpretation because of overlapping contributions from phenylalanine and other molecules¹⁸. Processed intensities at 1158 and 1518 cm⁻¹ were analyzed individually, and their equal-weight average was defined as the two-band score. Associations with β-carotene equivalent were summarized using Spearman rank correlations with 95% confidence intervals; Pearson correlation was also calculated as a secondary measure. Age- and sex-adjusted associations were calculated as correlations between residuals obtained by regressing rank-transformed variables on age and sex.

In the Raman PC1–PC2 plane, the carotenoid reference direction was defined by normalizing the sum of the loading vectors for the 1158 and 1518 cm⁻¹ variables. For each dietary variable, the standardized dietary value was regressed on PC1, PC2, age and sex, and the PC1/PC2 coefficients were normalized to define an association direction. Vector length was proportional to the incremental R² explained by PC1 and PC2 beyond the age/sex model, and cosine similarity with the carotenoid reference direction was calculated. In addition, Spearman correlations were calculated for all combinations of 70 BDHQ food variables and 549 Raman-shift variables to construct a descriptive Raman-shift × food correlation map.

### Statistical analysis and reproducibility

Continuous variables were summarized as mean (SD) or other appropriate descriptive statistics, and categorical variables as number and percentage. Trends in continuous variables across mAHEI-8 quartiles were assessed using Spearman tests, overall differences among quartiles using Kruskal–Wallis tests, and categorical variables using χ² tests. Sex differences in sodium intake were assessed using a two-sided Mann–Whitney U test, whereas age differences between participants with and without identified municipal records were assessed using the Kruskal–Wallis test. Wilson 95% confidence intervals were calculated for binomial proportions³⁴.

For Fig. 3b–d, healthcare-utilization analyses were restricted to the 129 participants with at least one identified medical claim and were performed without zero filling. Hospitalization was analyzed using logistic regression and total medical-care days using negative-binomial regression (NB2), with mAHEI-8 quartile entered as a categorical variable with Q1 as the reference and adjustment for age and sex. Because the Poisson model showed marked overdispersion, the negative-binomial model was used for total medical-care days. Benjamini–Hochberg correction was applied to the 28-variable cross-modal screen; for other exploratory/descriptive analyses, unadjusted statistics are reported as indicated in the text and figure legends.

Analyses were performed primarily in Python 3.12.8 (NumPy 1.26.4, pandas 2.2.2, SciPy 1.13.1, scikit-learn 1.5.1³⁵, statsmodels³⁶, umap-learn 0.5.7, PHATE 2.0.0 and matplotlib 3.9.2), with R 4.5.1 used for initial processing of municipal records. Fixed random seeds were used and recorded in the analysis code. This was an observational study using an existing cohort; no randomization or blinding was performed. Sample size was determined by participation in the FY2022 survey and availability of the respective measurements and was not based on an a priori power calculation.

### Use of generative artificial intelligence

Large language models, ChatGPT (OpenAI) and Claude (Anthropic), were used for manuscript structuring, English-language editing and programming support during data analysis. The analytical methods, code logic, numerical outputs, scientific interpretations and cited literature were independently verified by the authors, who take responsibility for the accuracy and completeness of the manuscript.

## Reporting summary

Further information on research design is available in the Nature Portfolio Reporting Summary linked to this article.

## Data availability

This study used participant-level health information from the Tsukuba Happiness Life Study linked at the individual level to medical-insurance claims and municipal health-check records held by Tsukuba City. Because these data contain personal health information and are subject to data-governance requirements of the municipality and related institutions, unrestricted public release of participant-level raw data is not permitted. De-identified data required to verify the study findings may be made available upon reasonable request to the corresponding author.

## Code availability

Analysis code used for data preprocessing, Raman spectral processing, dimensionality reduction, the cross-modal screen, multimodal prediction and figure generation is publicly available via figshare (DOI: 10.6084/m9.figshare.34004559). The repository includes code corresponding to the main and supplementary analyses and figures, together with input specifications, analytical populations, random seeds and major software versions.

## Acknowledgements

We thank Takahiro Yano (University of Tsukuba) for data management and Naomi Kaneko (University of Tsukuba) for her assistance with serum Raman measurements. We also thank Yasuko Maruyama, Naoko Onodera, and Mayuko Kano (University of Tsukuba), as well as Shoji Nakayama (National Institute for Environmental Studies), for their valuable support.

## Author contributions

K.H. and T.M. conceived and designed the study and jointly supervised the research. K.H., T.M., and T.N. performed the analyses. J.Y., K.S., S.N., X.Y., T.Shi, L.W., H.S., H.I., T.Sakurai, H.N., T.O., and M.S. contributed to the interpretation and critical evaluation of the analytical methods and results. K.H., T.M., and T.N. drafted the manuscript. All authors critically reviewed the manuscript, approved the final version, and agreed to be accountable for their respective contributions.

## References

1. Chiuve, S. E. et al. Alternative dietary indices both strongly predict risk of chronic disease. J. Nutr. 142, 1009–1018 (2012).

2. Tessier, A. J. et al. Optimal dietary patterns for healthy aging. Nat. Med. 31, 1644–1652 (2025).

3. Nohara, J. et al. Development of a concise healthy diet score for cardiovascular disease among Japanese: The Japan Collaborative Cohort Study. J. Atheroscler. Thromb. 31, 1443–1459 (2024).

4. Oono, F. et al. Associations of diet quality indices with all-cause and cause-specific mortality among Japanese adults in the Takayama study. Br. J. Nutr. 135, 737–748 (2026).

5. Zeevi, D. et al. Personalized nutrition by prediction of glycemic responses. Cell 163, 1079–1094 (2015).

6. Berry, S. E. et al. Human postprandial responses to food and potential for precision nutrition. Nat. Med. 26, 964–973 (2020).

7. Guillermo, C. et al. Diet quality and biomarker profiles related to chronic disease prevention: The Multiethnic Cohort Study. J. Am. Coll. Nutr. 39, 216–223 (2020).

8. Akbaraly, T. et al. Association of circulating metabolites with healthy diet and risk of cardiovascular disease: analysis of two cohort studies. Sci. Rep. 8, 8620 (2018).

9. Bagheri, M. et al. A lipid-related metabolomic pattern of diet quality. Am. J. Clin. Nutr. 112, 1613–1630 (2020).

10. Mayne, S. T. et al. Noninvasive assessment of dermal carotenoids as a biomarker of fruit and vegetable intake. Am. J. Clin. Nutr. 92, 794–800 (2010).

11. Ahn, S. et al. Validation of resonance Raman spectroscopy-measured skin carotenoid status as a biomarker for fruit and vegetable intake in Korean adults. Br. J. Nutr. 130, 1993–2001 (2023).

12. Cao, Y. et al. Sleep regularity is associated with cognitive function and shows an inverted U-shaped relationship with serum brain-derived neurotrophic factor. Sleep Med. 134, 106688 (2025).

13. Beltran Patino, V., et al. Associations of socioeconomic status and group exercise participation with happiness in older Japanese adults: A mediation analysis. Geriatr. Gerontol. Int. 26, e70625 (2026).

14. Pan, Z. et al. Associations between finger-dexterity dual task and neuropsychological tests in middle-aged and older adults. Ann. Geriatr. Med. Res. (2026). 10.4235/agmr.26.0086

15. Kobayashi, S. et al. Comparison of relative validity of food group intakes estimated by comprehensive and brief-type self-administered diet history questionnaires against 16-d dietary records in Japanese adults. Public Health Nutr. 14, 1200–1211 (2011).

16. Kobayashi, S. et al. Both comprehensive and brief self-administered diet history questionnaires satisfactorily rank nutrient intakes in Japanese adults. J. Epidemiol. 22, 151–159 (2012).

17. Udensi, J., Loughman, J., Loskutova, E. & Byrne, H. J. Raman spectroscopy of carotenoid compounds for clinical applications—a review. Molecules 27, 9017 (2022).

18. Suksuratin, P. et al. Rapid label-free detection of cholangiocarcinoma from human serum using Raman spectroscopy. PLoS ONE 17, e0275362 (2022).

19. Yu, D. et al. Adherence to dietary guidelines and mortality: a report from prospective cohort studies of 134,000 Chinese adults in urban Shanghai. Am. J. Clin. Nutr. 100, 693–700 (2014).

20. Oono, F. et al. Development of a Diet Quality Score for Japanese and comparison with existing diet quality scores regarding inadequacy of nutrient intake. J. Nutr. 153, 798–810 (2023).

21. Garcia-Perez, I. et al. Objective assessment of dietary patterns by use of metabolic phenotyping: a randomised, controlled, crossover trial. Lancet Diabetes Endocrinol. 5, 184–195 (2017).

22. Posma, J. M. et al. Nutriome–metabolome relationships provide insights into dietary intake and metabolism. Nat. Food 1, 426–436 (2020).

23. Zhu, K. et al. Proteomic signatures of healthy dietary patterns are associated with lower risks of major chronic diseases and mortality. Nat. Food 6, 47–57 (2025).

24. Radtke, M. D. et al. Criterion-related validity of spectroscopy-based skin carotenoid measurements as a proxy for fruit and vegetable intake: a systematic review. Adv. Nutr. 11, 1282–1299 (2020).

25. Colby, S. et al. Development and validation of the Short Healthy Eating Index Survey with a college population to assess dietary quality and intake. Nutrients 12, 2611 (2020).

26. Udensi, J., Loskutova, E., Loughman, J. & Byrne, H. J. Raman spectroscopic analysis of human blood serum of glaucoma patients supplemented with macular pigment carotenoids. J. Biophotonics 17, e202400060 (2024).

27. Kobayashi, M., Sasaki, S. & Tsugane, S. Validity of a self-administered food frequency questionnaire used in the 5-year follow-up survey of the JPHC Study Cohort I to assess carotenoids and vitamin C intake: comparison with dietary records and blood level. J. Epidemiol. 13, S82–S91 (2003).

28. Savitzky, A. & Golay, M. J. E. Smoothing and differentiation of data by simplified least squares procedures. Anal. Chem. 36, 1627–1639 (1964).

29. Jolliffe, I. T. & Cadima, J. Principal component analysis: a review and recent developments. Philos. Trans. A Math. Phys. Eng. Sci. 374, 20150202 (2016).

30. Moon, K. R. et al. Visualizing structure and transitions in high-dimensional biological data. Nat. Biotechnol. 37, 1482–1492 (2019).

31. McInnes, L., Healy, J. & Melville, J. UMAP: uniform manifold approximation and projection for dimension reduction. Preprint at https://arxiv.org/abs/1802.03426 (2018).

32. Benjamini, Y. & Hochberg, Y. Controlling the false discovery rate: a practical and powerful approach to multiple testing. J. R. Stat. Soc. B 57, 289–300 (1995).

33. Hoerl, A. E. & Kennard, R. W. Ridge regression: biased estimation for nonorthogonal problems. Technometrics 12, 55–67 (1970).

34. Wilson, E. B. Probable inference, the law of succession, and statistical inference. J. Am. Stat. Assoc. 22, 209–212 (1927).

35. Pedregosa, F. et al. Scikit-learn: machine learning in Python. J. Mach. Learn. Res. 12, 2825–2830 (2011).

36. Seabold, S. & Perktold, J. Statsmodels: econometric and statistical modeling with Python. Proc. 9th Python in Science Conf. 92–96 (2010).

