## Supplementary information for "Multimodal phenotyping reveals plant-food-related dietary information in serum Raman spectra of community-dwelling Japanese adults"

- Supplementary Table 1 | Correspondence between mAHEI-8 and the original AHEI-2010
- Supplementary Fig. 1 | Scoring characteristics of mAHEI-8 in this cohort
- Supplementary Fig. 2 | Presence of municipal records during 2014–2018 in relation to mAHEI-8 and age
- Supplementary Fig. 3 | Preceding health profile from Fig. 3a stratified by age and sex
- Supplementary Fig. 4 | PCA and UMAP representations of mAHEI-8 quartiles
- Supplementary Fig. 5 | Negative controls for the cross-modal screen
- Supplementary Fig. 6 | Diagnostics and sensitivity analyses for β-carotene-equivalent prediction
- Supplementary Fig. 7 | Raman shift × BDHQ food correlation map
- Supplementary Fig. 8 | Technical assessment of serum Raman peak-top flatness

**Supplementary figure legends**

**Supplementary Fig. 1 | Scoring characteristics of mAHEI-8 in this cohort.**

**a**, Scoring functions and intake distributions in this cohort for the seven components scored using fixed thresholds. **b**, Component-score distributions for all eight components. **c**, Correspondence between sex-specific sodium-intake deciles and sodium component scores.

**Supplementary Fig. 2 | Presence of municipal records during 2014–2018 in relation to mAHEI-8 and age.**

**a**, Proportion of participants with at least one identified municipal record in each mAHEI-8 quartile (Q1–Q4: 32/101, 41/100, 31/100 and 28/100; χ² P = 0.232). **b**, Age distributions of the 132 participants with at least one identified municipal record and the remaining 269 participants. Participants with records were older than those without records.

**Supplementary Fig. 3 | Preceding health profile from Fig. 3a stratified by age and sex.**

The same variables, claims handling and row order as in Fig. 3a were applied to four strata: men <65 years, women <65 years, men ≥65 years and women ≥65 years. Each variable was row-standardized within each stratum; colors should therefore be interpreted only as the quartile pattern within each panel. Cells without an available health-check measurement were left blank.

**Supplementary Fig. 4 | PCA and UMAP representations of mAHEI-8 quartiles.**

PCA and UMAP representations of the same four phenotypic spaces as in Fig. 4a were constructed among the 370 participants with complete blood/urine, serum metabolome and quality-controlled Raman data. The 367 participants with corresponding mAHEI-8 data are shown and colored by mAHEI-8 quartile. Dietary variables, participant identifiers, visit date, age and sex were not used to construct the representations.

**Supplementary Fig. 5 | Negative controls for the cross-modal screen.**

Ten real dietary outcomes were randomly shuffled across participants to generate negative-control outcomes and were analyzed using the same cross-validation and Raman-block permutation procedures as in Fig. 5a. **a**, Raman incremental OOF R² for the ten shuffled outcomes. Filled and open circles indicate models with blood + metabolome and blood + metabolome + age + sex as the baseline, respectively. Incremental OOF R² values were distributed around zero, and no negative control had permutation P < 0.05. **b**, Calibration of permutation P values for the shuffled outcomes. Observed permutation P values are plotted against the quantiles expected under a uniform distribution; agreement with the diagonal indicates the behavior expected under the null. Negative controls were not included in the 28-outcome Benjamini–Hochberg correction.

**Supplementary Fig. 6 | Diagnostics and sensitivity analyses for β-carotene-equivalent prediction.**

**a**, OOF residuals versus predictions from the three-layer model. **b**, Normal Q–Q plot of OOF residuals. **c**, Observed-versus-predicted plot for the blood + metabolome model without Raman. **d**, OOF R² for models with and without visit date added to all model combinations. **e**, Raman incremental OOF R² on the raw β-carotene-equivalent scale and after log(1+x) transformation. Error bars in **d,e** show the s.d. across 10 cross-validation repeats.

**Supplementary Fig. 7 | Raman shift × BDHQ food correlation map.**

Among 370 participants with quality-controlled Raman spectra and BDHQ data, Spearman correlations were calculated for 38,430 combinations of 70 food variables × 549 Raman-shift variables. Dashed lines indicate 1158 and 1518 cm⁻¹, the regions interpreted molecularly in Fig. 6. Because each Raman shift can contain contributions from multiple molecules and the food variables are themselves correlated, no molecular assignment was applied to individual cells.

**Supplementary Fig. 8 | Technical assessment of serum Raman peak-top flatness.**

**a**, Representative raw spectra with flat-topped peaks around carotenoid-associated bands. **b**, Maximum raw detector counts across all 376 spectra. The three spectra excluded during technical QC are indicated. Maximum counts did not show a clear common upper limit across spectra. **c**, Total raw intensity, local background at 1158 cm⁻¹ and background-subtracted carotenoid-band height for spectra flagged for flatness, shown as cohort percentiles. **d**, Application of a common plateau-width classification rule to the instrument-native axis, 2 cm⁻¹ common axis and primary preprocessed matrix. Classification was affected by the Raman-shift grid and was largely resolved after preprocessing. The primary matrix (n = 373) excludes the three spectra removed by technical QC; no additional spectra were excluded on the basis of peak-top flatness.

**Supplementary table legend**

**Supplementary Table 1 | Correspondence between mAHEI-8 and the original AHEI-2010.**

Of the 11 original AHEI-2010 components, vegetables, fruit, long-chain omega-3 fatty acids, PUFA, red + processed meat and sodium were retained. Nuts + legumes was represented by legumes because nuts could not be independently quantified. Sugar-sweetened beverages + fruit juice was calculated from cola and 100% fruit juice. Whole grains, trans fat and alcohol were excluded. Sodium was scored using sex-specific intake deciles in this cohort. Each component was scored from 0 to 10, giving a total mAHEI-8 range of 0–80. Serving definitions and component-specific scoring thresholds are provided in the table.
