## Supplementary figures for "Multimodal phenotyping reveals plant-food-related dietary information in serum Raman spectra of community-dwelling Japanese adults"

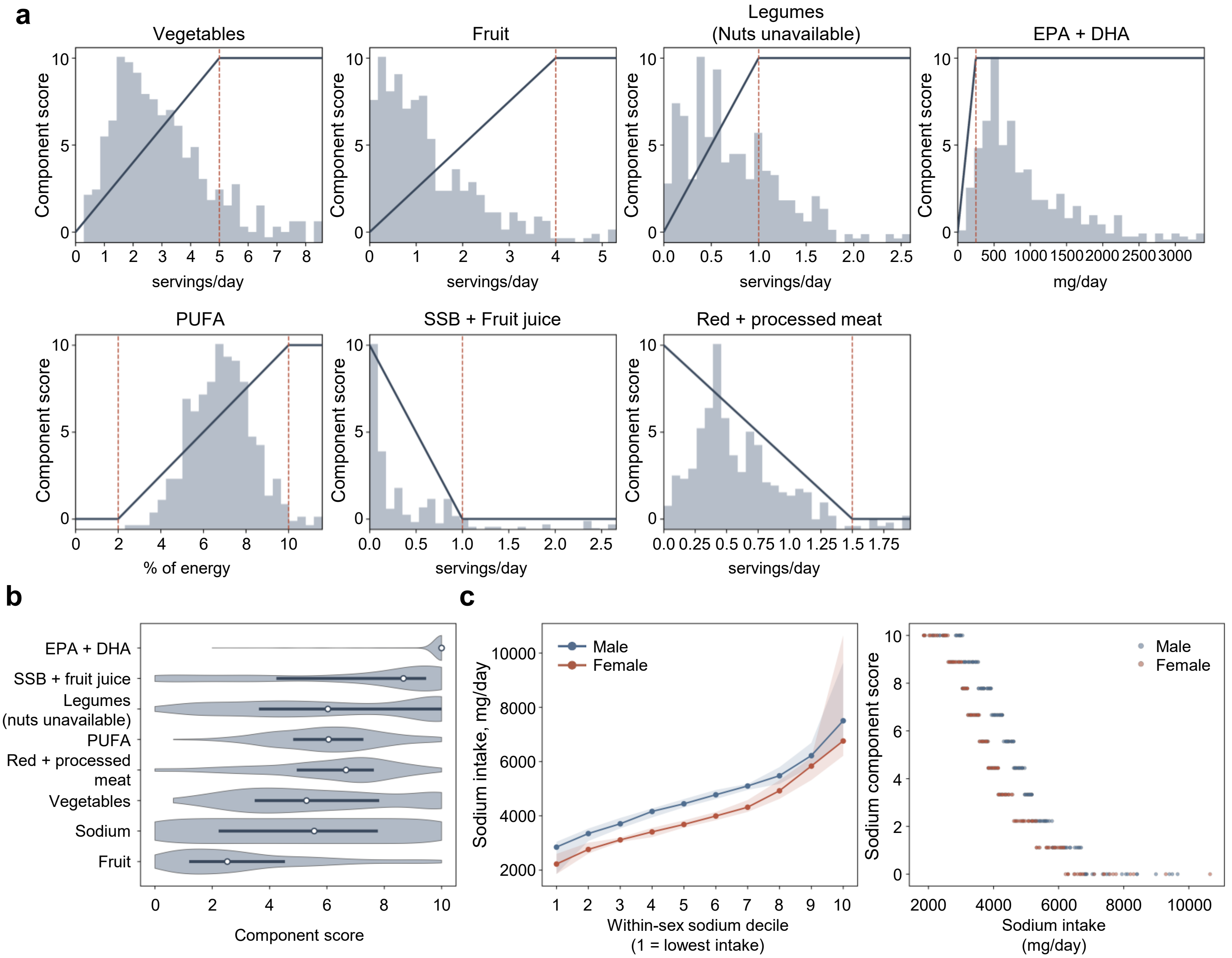


Supplementary Figure 1. Hiraga et al.

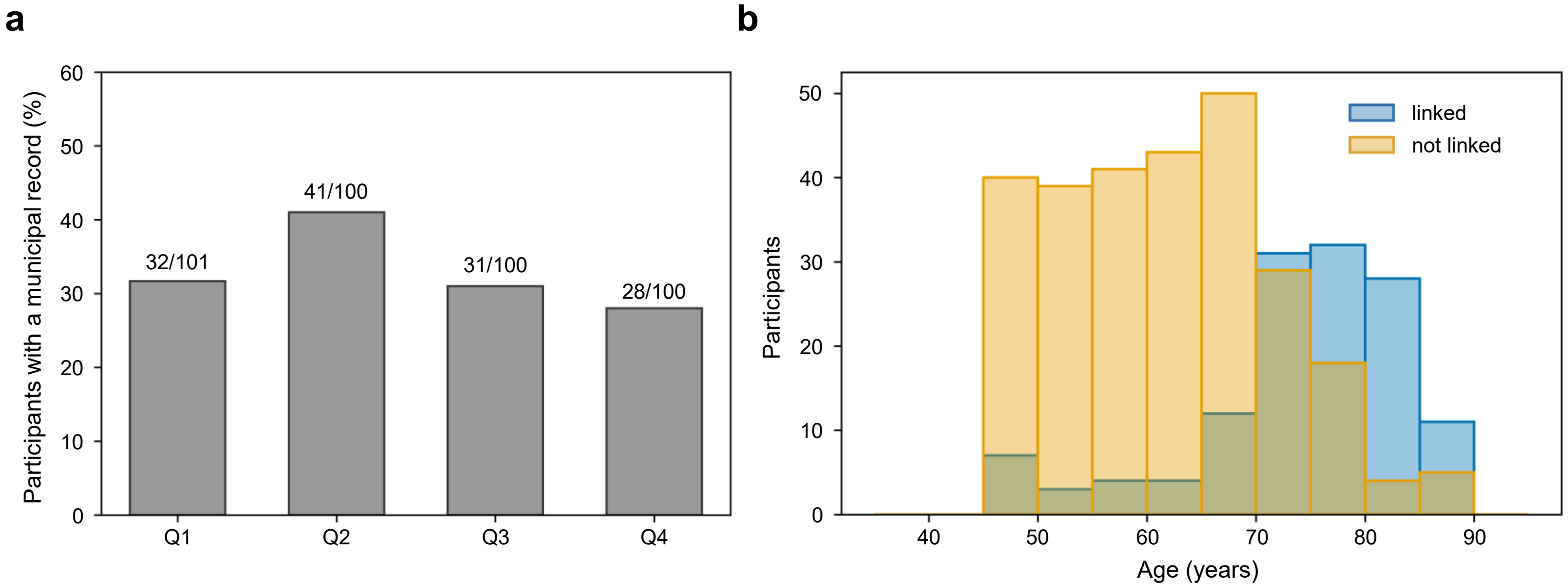


Supplementary Figure 2. Hiraga et al.

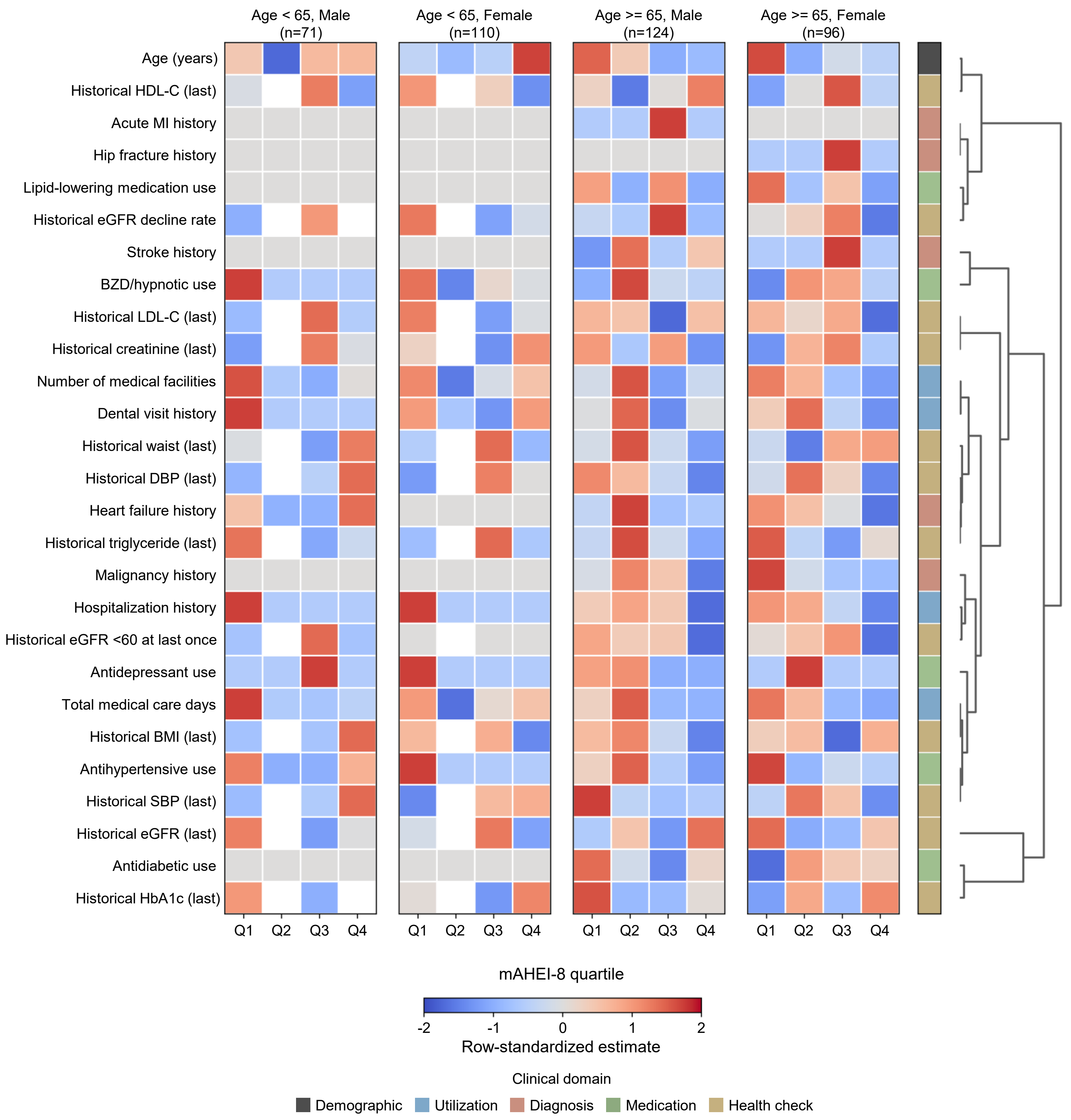


Supplementary Figure 3. Hiraga et al.

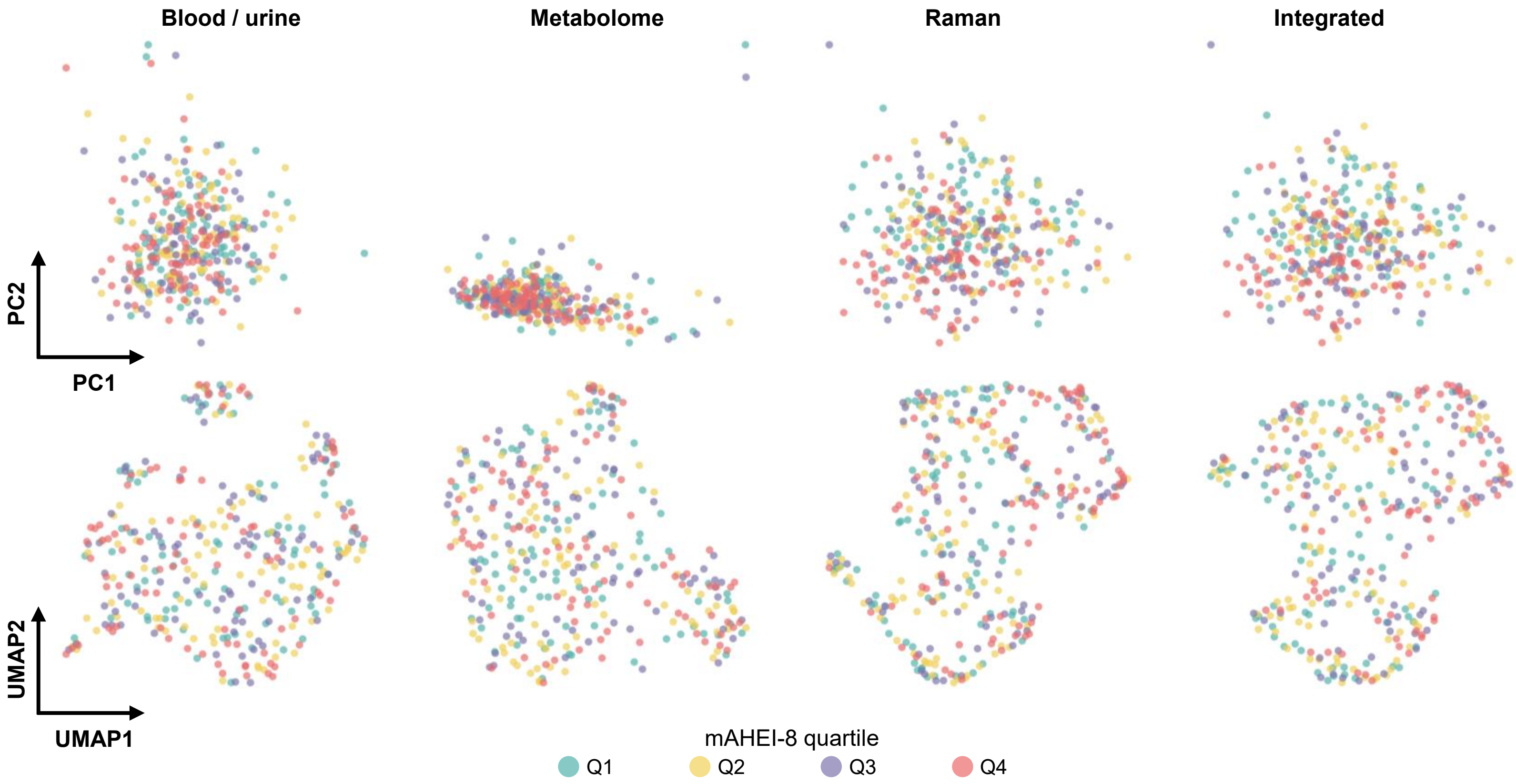


Supplementary Figure 4. Hiraga et al.

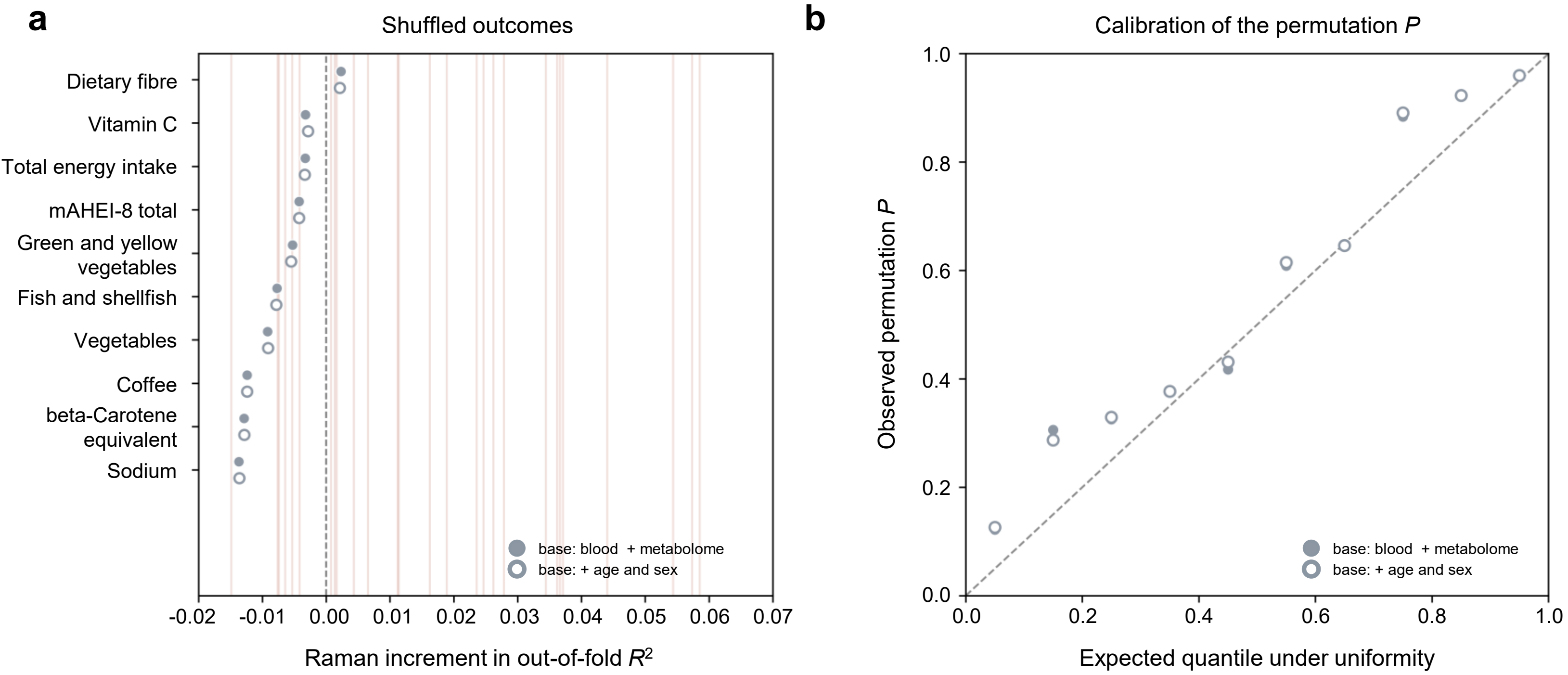


Supplementary Figure 5. Hiraga et al.

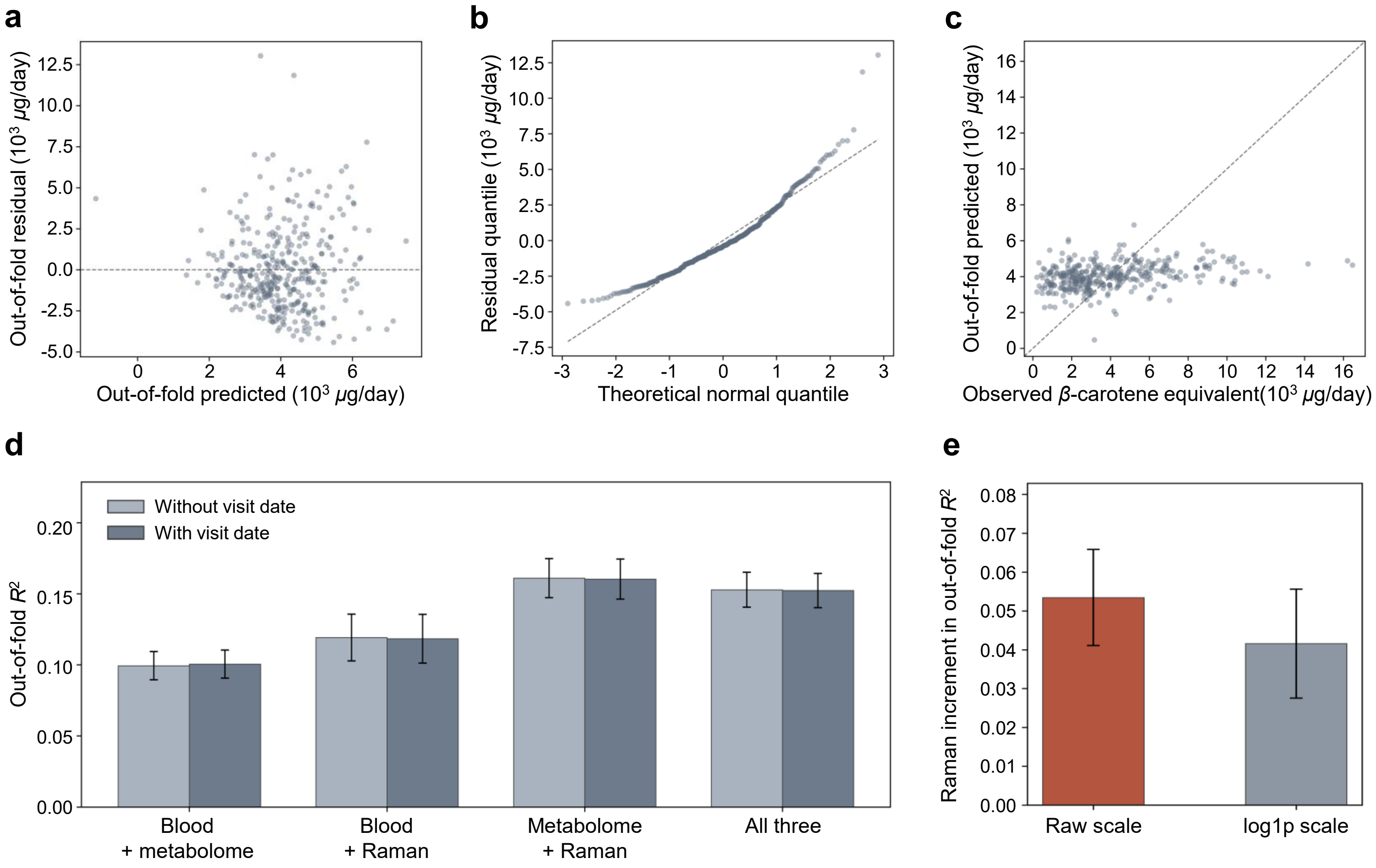


Supplementary Figure 6. Hiraga et al.

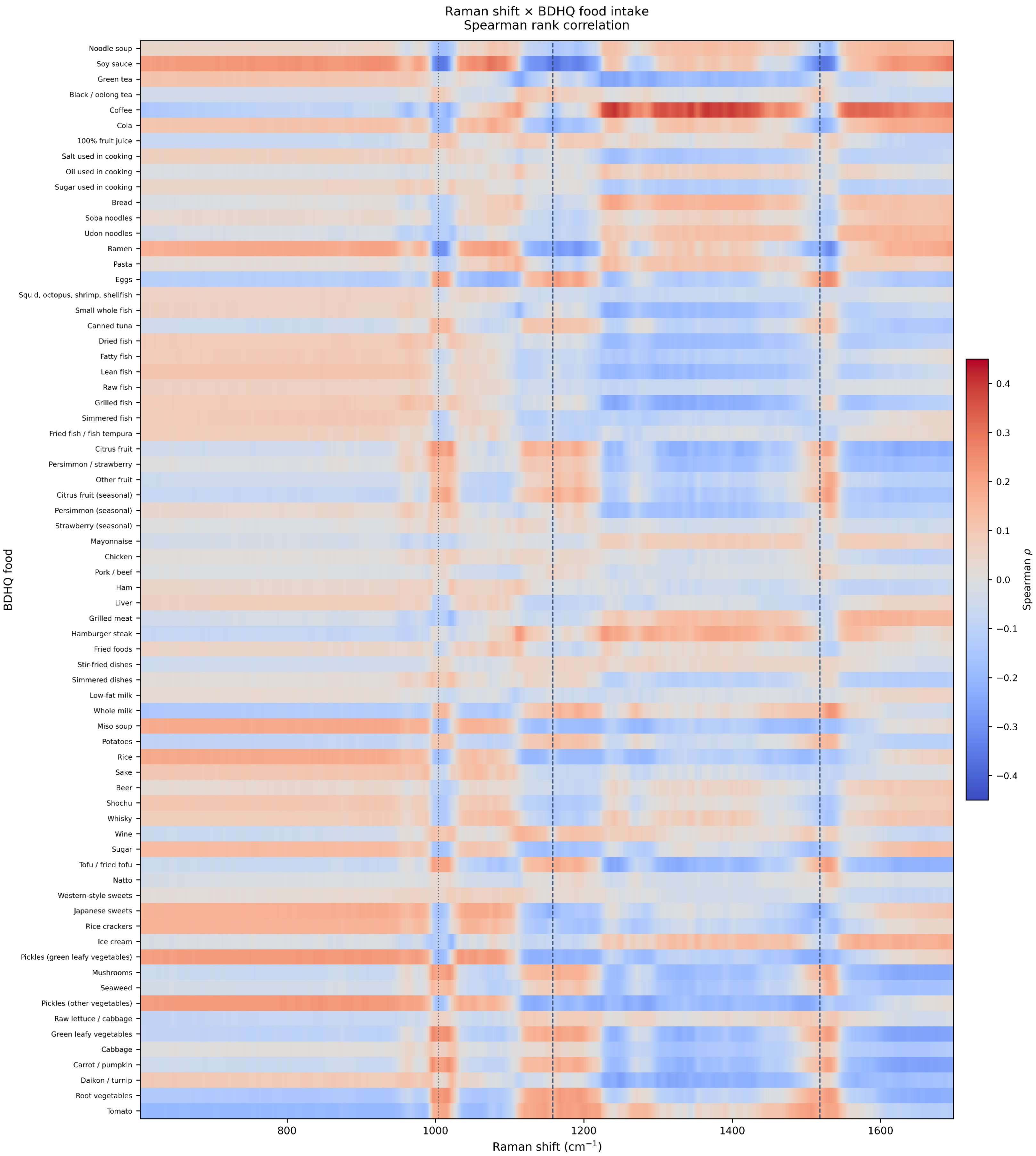


Supplementary Figure 7. Hiraga et al.

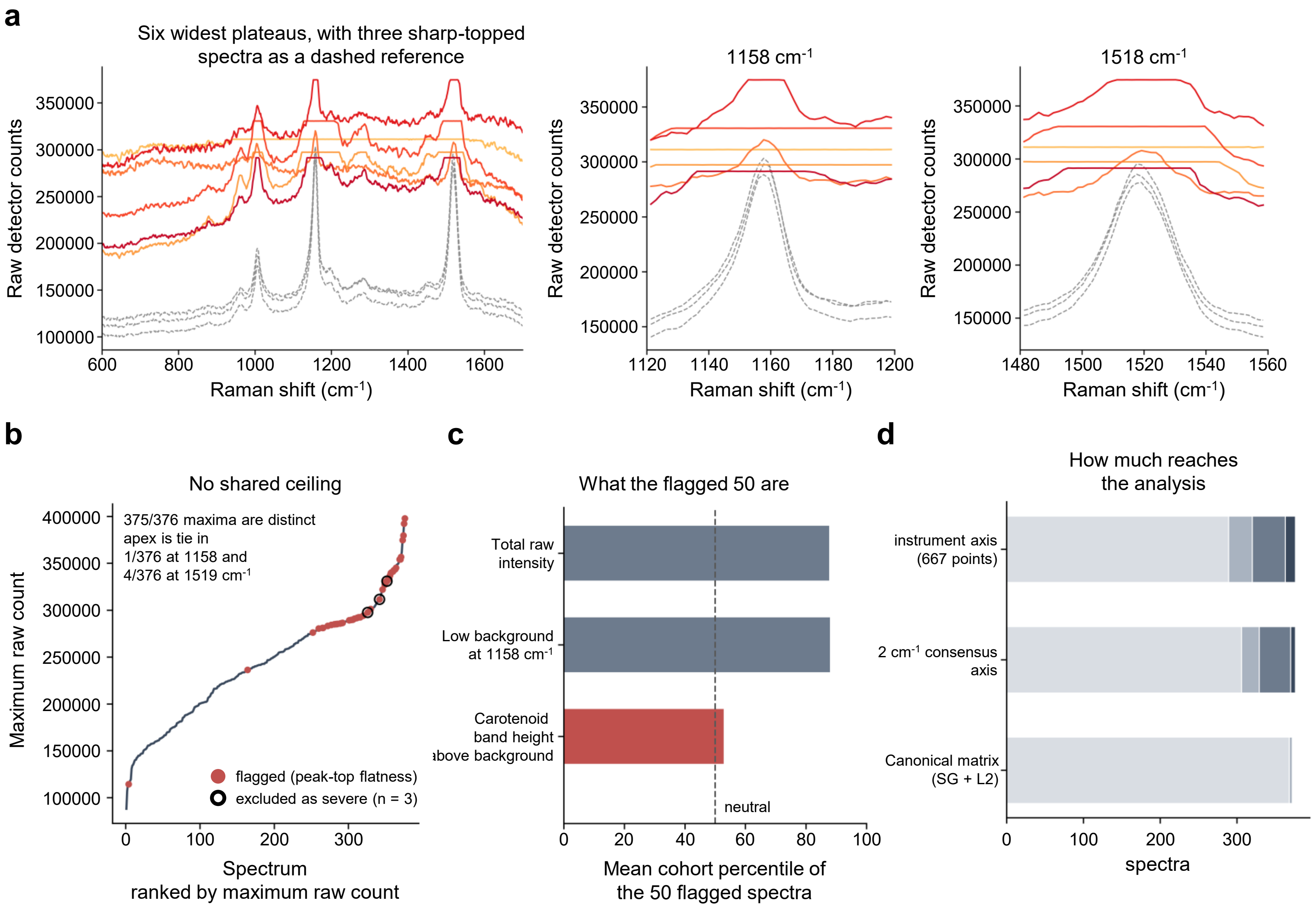


Supplementary Figure 8. Hiraga et al.

| AHEI-2010 component | Status | mAHEI-8 component | BDHQ source | Serving_definition | Scoring rule |
|---|---|---|---|---|---|
| Vegetables | retained | Vegetables | f_nutr1: W_GV + W_OV | 100 g per serving | 0 at 0 servings/day, 10 at >= 5 servings/day, linear between |
| Fruit | retained | Fruit | f_nutr1: W_FR | 100 g per serving | 0 at 0 servings/day, 10 at >= 4 servings/day, linear between |
| Whole grains | omitted | *not included* | | | not derivable from the available BDHQ variables with a consistent definition |
| Nuts and legumes | adapted | Legumes | f_nutr1: W_PL | 100 g per serving | 0 at 0 servings/day, 10 at >= 1 serving/day, linear between |
| Long-chain omega-3 fatty acids | retained | EPA + DHA | nutr: C205N3 + C226N3 | mg per day | 0 at 0 mg/day, 10 at >= 250 mg/day, linear between |
| Polyunsaturated fatty acids | retained | PUFA | nutr: PFA and EN | percentage of total energy | 0 at <= 2 % energy, 10 at >= 10 % energy, linear between |
| Sugar-sweetened beverages and fruit juice | adapted | SSB + fruit juice | food: BEV4 (cola) + BEV5 (100 % juice) | 248 g per serving | 10 at 0 servings/day, 0 at >= 1 serving/day, linear between |
| Red and processed meat | retained | Red and processed meat | food: MEAT2 (pork and beef), MEAT3 (ham) | 100 g unprocessed, 50 g processed | 10 at 0 servings/day, 0 at >= 1.5 servings/day, linear between |
| Trans fat | omitted | *not included* | | | not derivable from the available BDHQ variables |
| Sodium | retained | Sodium | nutr: NA | within-sex deciles of intake | 10 for the lowest decile, 0 for the highest, linear between |
| Alcohol | omitted | *not included* | | | not included in the study-specific score |

Supplementary Table 1. Hiraga et al.
